# The tortured past of young polymorphic sex chromosomes revealed through multiple *de novo* genome assemblies of the mountain pine beetle

**DOI:** 10.64898/2026.01.27.702068

**Authors:** German Lagunas-Robles, Barbara J. Bentz, Medhavi Verma, Jim Vandygriff, Matt Hansen, Tyler Simmonds, Renee Corpuz, Angela Kauwe, Scott Geib, Ryan R. Bracewell

## Abstract

Neo-sex chromosomes provide a powerful system for studying the early stages of sex chromosome evolution and the genomic mechanisms that may contribute to reproductive isolation. Using PacBio long-read HiFi sequencing, Hi-C scaffolding, and sex-specific transcriptomic data, we generated six chromosome-level assemblies (male and female from three populations) of the mountain pine beetle (*Dendroctonus ponderosae*), a species known to harbor three partially reproductively isolated neo-Y haplogroups. These assemblies reveal that the large neo-X and neo-Y chromosomes formed through sequential fusions of the ancestral X with three autosomes, with recombination cessation occurring at ∼8.6, ∼6.3, and ∼4.3 MYA for each event. Comparative analyses show that while neo-X chromosomes remain largely collinear across populations, neo-Ys exhibit dramatic structural divergence, with 900–1,200 inverted segments per haplogroup and only ∼65% of sequence able to be aligned to the neo-X. Repeat analyses demonstrate moderate TE accumulation on the neo-Y, particularly LTR elements, and gene mapping analyses reveal extensive degeneration: ∼62% of neo-Y genes exhibit gene loss, fragmentation, or disruptive mutations. All populations retain a single pseudoautosomal region (PAR), though PAR size and gene content vary due to neo-Y specific rearrangements. Across neo-Ys, 27 genes are uniquely missing in the Western haplogroup, including previously identified candidates implicated in hybrid male sterility. Broader comparisons among neo-Ys show widespread structural variation, population specific patterns of degeneration, and limited gene family expansions. Together, these results provide the first full characterization of neo-sex chromosome evolution in *D. ponderosae*, revealing rapid, lineage specific neo-Y degeneration and highlighting the potential for sex chromosome divergence to contribute to emerging reproductive incompatibilities within a single species.

## INTRODUCTION

Sex chromosomes have captivated biologists since they were first discovered. These chromosomes arise when autosomes acquire sex-determining roles, often triggering genomic changes that ultimately produce highly heteromorphic chromosomes with major differences in gene content and regulation [1–3]. To understand how these processes unfold over millions of years, studies often take advantage of evolutionarily recent sex chromosome-autosome fusions, commonly referred to as neo-sex chromosomes (**Fig. 1A**). These studies reveal several generalities; following the loss of recombination between neo-sex chromosomes, the non-recombining chromosome (i.e., the Y or W) loses protein coding genes via degeneration, transposable elements proliferate, and the chromosome first expands, and then contracts in physical size, as the Y (or W) ages [4–6]. Interestingly, even in systems with fairly young neo-sex chromosomes, these neo-Y (or neo-W) changes appear to happen rather quickly after the loss of recombination, likely the result of the reduced effective population size of the sex-limited chromosome, and via linked selection and repeated selective sweeps [7, 8]. However, relatively little attention has been given to exploring how young sex chromosomes, and specifically the neo-Y (or neo-W) could be a hotbed for genomic changes that could segregate within a species and directly lead to the evolution of reproductive incompatibilities [9, 10].

**Figure 1.**
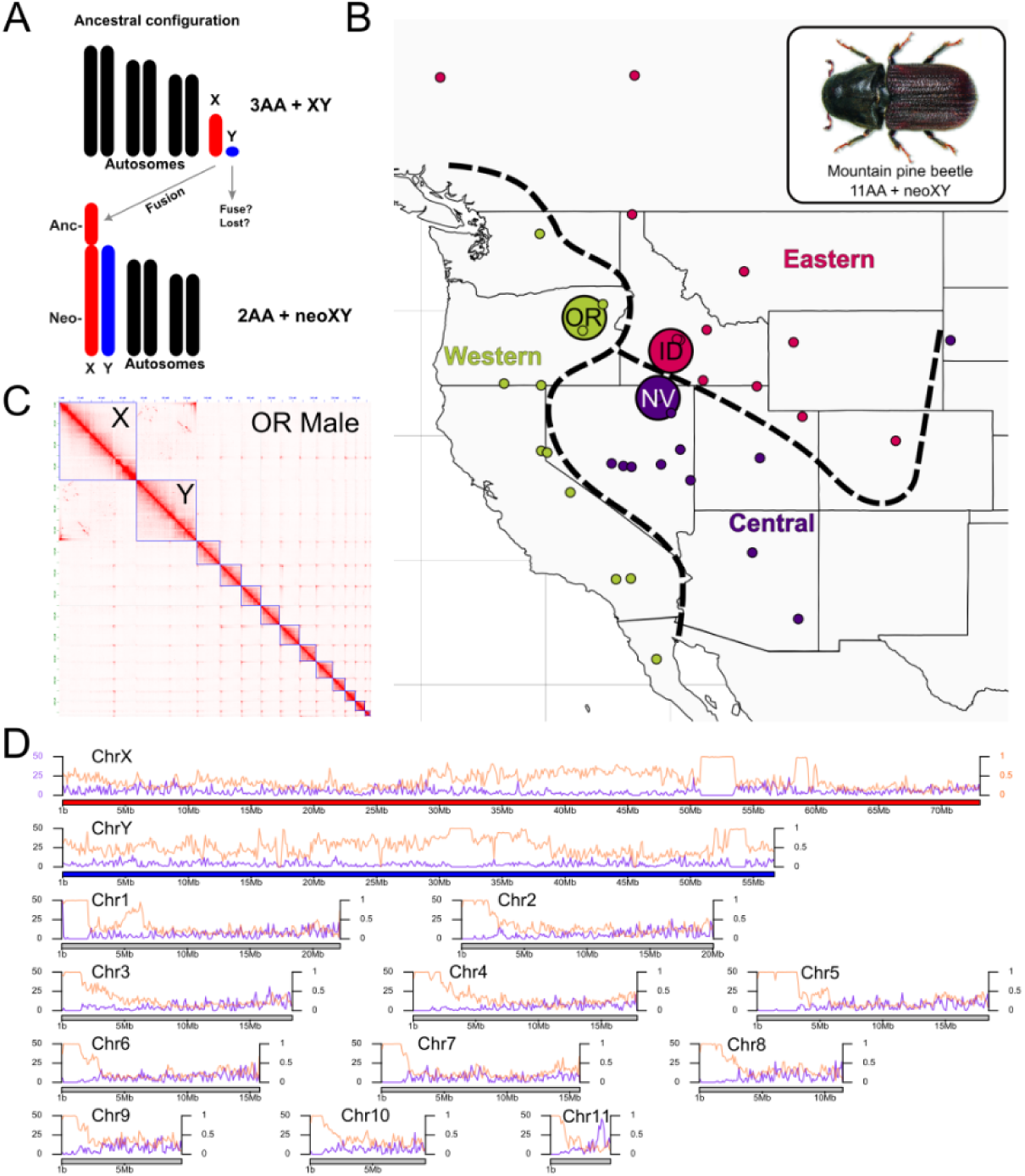
Six denovo chromosome-level assemblies of male and female mountain pine beetles reveal large sex chromosomes and complete neo-Y assembly. **A)** Neo-sex chromosomes are formed when ancestral sex chromosomes (often heteromorphic) fuse to autosomes. Shown is an example where an ancestral X (anc-X) fuses to an autosome resulting in a large X chromosome comprised of an anc-X and neo-X region, and a neo-Y chromosome. **B)** The mountain pine beetle (*Dendroctonus ponderosae*) is a major forest pest in western North America, and has 11 autosomes and large neo-XY sex chromosomes thought to be the result of an X-autosome fusion [26]. Previous studies have uncovered three distinct Y-haplogroups, denoted as Western, Eastern, and Central, with variable barriers to genetic exchange (hybrid sterility and inviability) in crosses between Western and Eastern/Central Y-haplogroups. Small dots represent Y-haplogroup status for various populations across the range of the species [12] and dashed lines delinieate putative geographic boundaries between Y-haplogroups. Large circles represent sample locations for this study. **C)** Hi-C contact map for a representative OR male assembly. Blue boxes denote chromosome boundaries. **D)** Genomic features of a the OR male chromosome-level genome assembly. Number of protein coding genes and proportion of sequence masked for repetitive sequences in 50 kb windows in purple (left axis) and orange (right axis), respectively. Sex chromosomes are shown in red (X) and blue (Y) while autosomes are shown in grey.

The mountain pine beetle is a unique system to investigate neo-sex chromosomes and how neo-Y evolution and degeneration could be important in the formation of reproductive incompatibilities, and thus, new species [11–13]. Previous work has shown that this important forest pest is comprised of three Y-haplogroups that vary in the degree of postzygotic reproductive incompatibilities in genetic crosses (**Fig. 1B**). Population genomic analyses suggest gene flow occurs between populations and across most genomic compartments (autosomes, X, mitochondria), yet the (neo)Y shows distinct geographic boundaries [11, 12]. These boundaries align with results from crosses that show Eastern and Central populations produce hybrid males with varying degrees of sterility and hybrid inviability when crossed to Western [11] (**Fig. 1B**); phenotypes well known to be associated with the earliest stages of speciation [14, 15]. Hybrid sterility has not been detected in crosses between Central and Eastern [16, 17], yet there is an atypical absence of recombination over a large segment of the neo-X chromosome in surviving males, suggestive of some hybrid male inviability/lethality [17]. Y-haplogroup specific gene loss (degeneration) has been proposed as a driver of these incompatibilities as hybrids would have unique neo-X/neo-Y gene combinations [11].

Although MPB was just the second beetle to have its genome assembled [18] and there is currently one chromosome-level assembly [19], the neo-Y has remained a mystery. Further, although sequencing breakthroughs are rapidly increasing the number of high-quality genome assemblies and the exploration of the non-recombining neo-sex chromosome [5, 8, 20–25], nearly all neo-sex chromosome study systems have yet to enter the pangenomics realm and intraspecific structural variation remains challenging to characterize and is largely unexplored. Here, we present multiple chromosome-level assemblies of MPB and include a full characterization of the neo-X and neo-Y, including populations known to have reproductive incompatibilities correlated with sex chromosome variation. We use long-read sequencing, Hi-C, and sex-specific annotations, to explore the evolutionary history of the neo-sex chromosomes and reveal an enormous number of structural differences between not only the neo-X and neo-Y, but also between three neo-Ys. Further, we detail methods for exploring neo-Y degeneration in a population context by using assembly-specific annotations coupled with gene lifting between neo-Ys to catalog gene presence/absence and the extent of pseudogenization.

## RESULTS/DISCUSSION

### Six *de novo* chromosome-level genome assemblies

To assemble chromosome-level genomes we used sex and population specific PacBio HiFi coupled with Hi-C to generate independent chromosome scale assemblies for one male and female MPB from three distinct populations representing the three previously identified Y-haplogroups [11, 12]. Beetle populations were collected from Oregon, Idaho, and Nevada and are hereafter abbreviated as OR, ID, and NV (**Fig. 1B**). We long-read sequenced males and females to median autosomal depths of 97× and 91×, respectively. The karyotypic formula for MPB is 11AA + neoXY and we unambiguously assembled 11 autosomes and the X and Y. The three male genome assemblies ranged from 270.7 – 293.2 Mb in total length (haploid autosomes + XY) while the female assemblies ranged from 219.0 – 237.6 Mb (haploid autosomes + X) (**sTable 1**). All assemblies were found to be of high quality, with BUSCO completeness ranging from 98.6 - 98.8 (**sTable 2**). Most autosomes assembled in a single contig and the sex chromosomes in just a few contigs that were unambiguously scaffolded using Hi-C (**Fig. 1C and sFig. 1**). To confirm the quality of the X and Y assembly, we first re-mapped the PacBio HiFi data to its respective assembly to confirm equal sequencing coverage over each chromosome (**sFig. 2**). We also mapped published male and female Illumina data to each male assembly and confirmed that females had double the coverage over the X and that the Y had predominantly male coverage (**sFig. 3**).

### Gene and repeat content

Transposable elements (TEs) and other repetitive sequences can make up a large fraction of a genome and are often highly enriched on sex chromosomes [27]. To fully characterize the repeat landscape of each assembly and begin to explore the sex chromosomes, we created six assembly-specific repeat libraries and identified and masked repetitive sequences prior to downstream gene annotation. On average, male assemblies were found to be 36.8% repetitive while female assemblies were 32.3% (**Table 1**). As expected, repetitive elements were not evenly distributed across each chromosome, and we found enrichment at one end of all autosomes suggestive of pericentromeric regions and either acro- or telocentric chromosomes (**Fig. 1D** and **sFig. 4**). The X and Y showed distinct enrichment elsewhere along each chromosome consistent with meta- or submetacentric chromosomes (**Fig. 1D and sFig. 4**). The majority of identifiable TEs in the MPB genome were LINEs (∼5.7% of masked bases), followed by DNA transposons (∼5%), and LTR elements (∼3.6%), although many of the repetitive elements remain unclassified and make up a substantial proportion of the genome (∼20%).

**Table 1.**
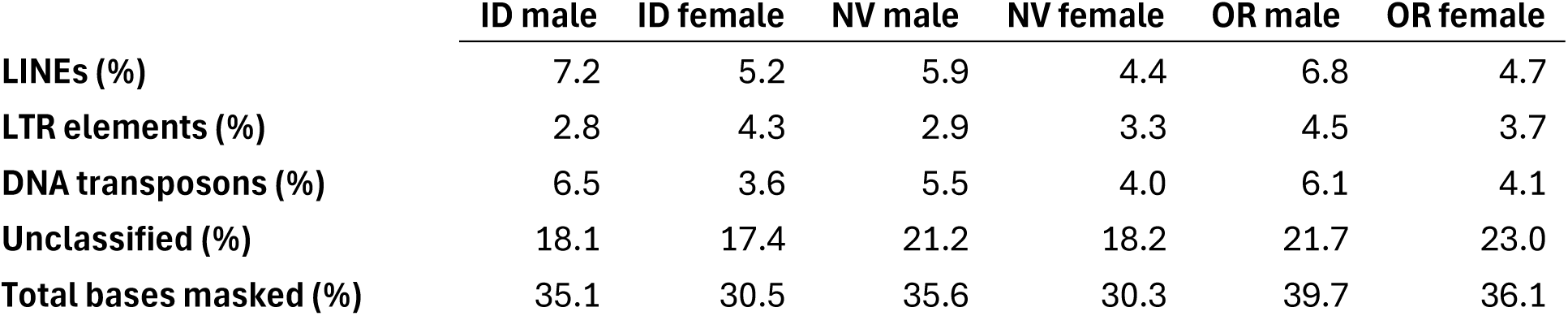
Repetitive element classes identified by RepeatModeler for each genome assembly.

|  | ID male | ID female | NV male | NV female | OR male | OR female |
| --- | --- | --- | --- | --- | --- | --- |
| **LINEs (%)** | 7.2 | 5.2 | 5.9 | 4.4 | 6.8 | 4.7 |
| **LTR elements (%)** | 2.8 | 4.3 | 2.9 | 3.3 | 4.5 | 3.7 |
| **DNA transposons (%)** | 6.5 | 3.6 | 5.5 | 4.0 | 6.1 | 4.1 |
| **Unclassified (%)** | 18.1 | 17.4 | 21.2 | 18.2 | 21.7 | 23.0 |
| **Total bases masked (%)** | 35.1 | 30.5 | 35.6 | 30.3 | 39.7 | 36.1 |

To identify protein-coding genes, we independently annotated each assembly to uncover unique genes and increase our confidence in sex chromosome annotation which is complicated by underlying sequence similarity between the neo-X and neo-Y in males. We used the Braker3 pipeline [28] with newly generated tissue-specific RNAseq data from male and female heads, ovaries and testes, supplemented with sex-specific predicted proteins from previous assemblies [19, 29]. Our female assemblies (autosomes + X) had an average of 12,313 protein coding genes while our male assemblies (autosomes + XY) had 14,542 (**sTable 3**). The X chromosome, which is the largest chromosome in the genome, was found to harbor ∼27% of all genes in a female assembly. These initial annotations suggested large and gene-rich neo-Ys (∼2k genes). As seen with repetitive elements, protein coding genes were not equally distributed across each chromosome. Regions with higher gene density were in regions with low repeat density (**Fig. 1D and sFig. 4**), a pattern observed in many animal genomes.

### Autosome and X Chromosome Synteny

We first explored synteny (chromosome conservation) and collinearity (gene order) between our six reference genomes using Genespace [30]. As expected, all chromosomes were syntenic, and 8 of 11 autosomes were found to be largely collinear (**Fig. 2A, sFig. 5**). However, three autosomes (Chr2, Chr3, Chr8) had at least one inversion distal the end enriched for repetitive elements (**Fig. 2A, sFig. 5**), and these ranged in size from 0.95 Mb to 6.7 Mb in length (**sTable 4**), with some complex rearrangements also being observed (e.g., OR male Chr3). The extent to which these inversions segregate broadly in natural populations is currently unknown, but several have been previously identified [17, 19]. Exploration of alternative haplotypes in our assemblies suggested several individuals were heterozygous for some inversions. Large segregating inversions have been found to be widespread in many systems [31] and may even be of adaptive importance [32] and play a role in speciation [33]. Interestingly, one small, but complex set of inversions was detected on the X chromosome in a region also found enriched for repetitive elements, and OR, ID, and NV each had a unique configuration of the region (**Fig. 2A**).

**Figure 2.**
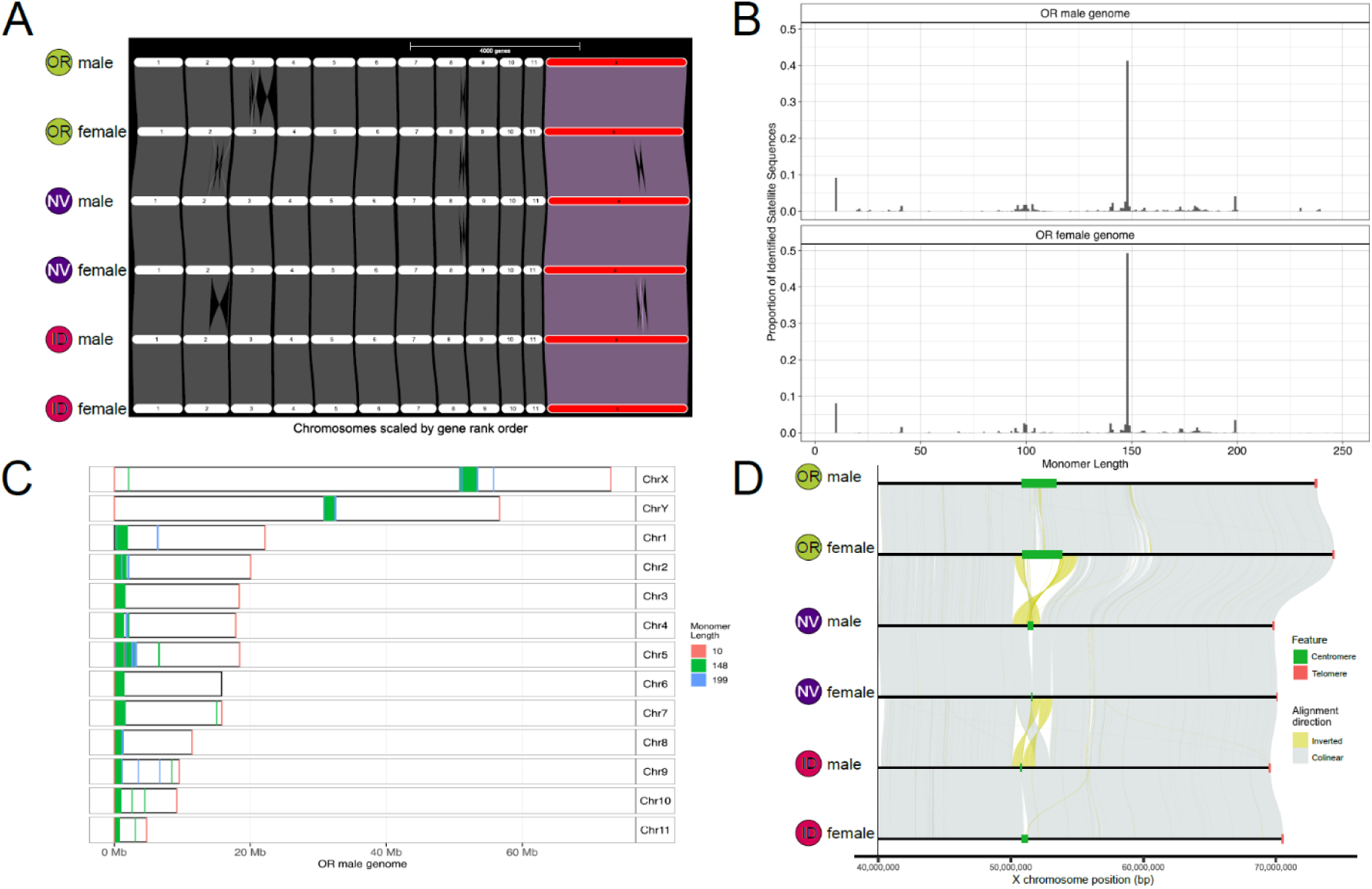
Synteny analysis and identification of putative centromeric/telomeric satellite sequences uncover chromosomal structure. **A)** Autosome and X chromosome synteny between all six chromosome-level genome assemblies. X chromosomes shown in red with collinear regions connected via purple ribbons. Note overall autosomal and X chromosome conservation and several autosomal inversions. **B)** Proportion of satellite sequences identitified in OR male and female genomes. Peaks represent the putative centromere monomers (148-mer and 199-mer) and telomeric monomer (10-mer). (see **sFig. 6** for other assemblies). **C)** Chromosomal location of putative centromeric and telomeric satellite sequence in reference OR male genome (see **sFig. 7** for other assemblies). **D)** Fine-scale whole genome alignment between six X chromosomes in a region with Y-haplogroup-specific inversions (yellow ribbons). Centromere location shown in green and telomere location in orange. Note the putative centromere was a universal breakpoint in all assemblies and the high coverage OR male and female assemblies traversed furthest into this region (see **sFig. 2**).

### Identification of putative centromeric and telomeric satellite sequence

Centromeres and telomeres are often comprised of long arrays of satellite sequences, and in some beetles species, these sequences have been identified [34–36]. Our TE analyses suggested centromere location varied for autosomes and sex chromosomes and therefore we set out to identify the putative centromere-associated satellite and telomeric sequence using a combination of TRASH [37], ModDotPlot [38], and manual inspection. We found three motifs to be highly abundant in all genome assemblies (**Fig. 2B** and **sFig. 6**). We found a short telomeric motif, TTAGTTTGGG, and long arrays of this sequence were found on the ends of our assembled chromosomes (**Fig. 2B, sFig. 7, sTable 5**). We also found two putative centromere-associated satellite sequences; a 148 bp and 199 bp motif, with underlying sequence similarity between the two, suggesting one is a complex of the other (**Fig. 2C**, **sFig. 7**, and **sTable** 5). These satellite sequences were highly enriched in our assemblies, although varying in total assembled length depending on the initial sequencing depth and overall assembly quality. These sequences were localized to the end of all autosomes yet were found in more central locations on the X and Y chromosome (**Fig. 2C, sTable 6**). Interestingly, the complex of inversions found on the X flank these centromere-associated satellite sequences, suggesting complex peri- and paracentric inversions separate alternative X configurations found in our NV, ID, and OR individuals (**Fig. 2D**). Although the functional significance of these centromere-associated inversions among these populations is unknown, crosses between an Eastern and Central population have shown suppressed recombination in this region [17].

### What fused to create the neo-sex chromosomes?

Given the large size of the X and Y chromosome, we next aimed to clarify the evolutionary events and putative fusion(s) between autosomes and the ancestral-X (anc-X) that gave rise to the neo-sex chromosomes in MPB. We compared our ID female assembly to a chromosome-level assembly of *D. valens* [39], a species considered to harbor a more ancestral *Dendroctonus* karyotype comprised of 13 autosomes and typical beetle XY sex chromosomes [40]. Using 6,902 1:1 orthologs, we found most autosomes are conserved and syntenic, with only one fusion/fission event between the two species (**Fig. 3A**). In contrast, the MPB X chromosome appeared to be a composite of four separate *D. valens* chromosomes (**Fig. 3A**). This composite MPB X chromosome featured the ancestral-X (anc-X) on one end, with three separate *D. valens* autosomes (Chr1, Chr4, Chr11) mapping to other regions, demonstrating at least one, and suggesting potentially several X-autosomal fusions (**Fig. 3A**).

**Figure 3.**
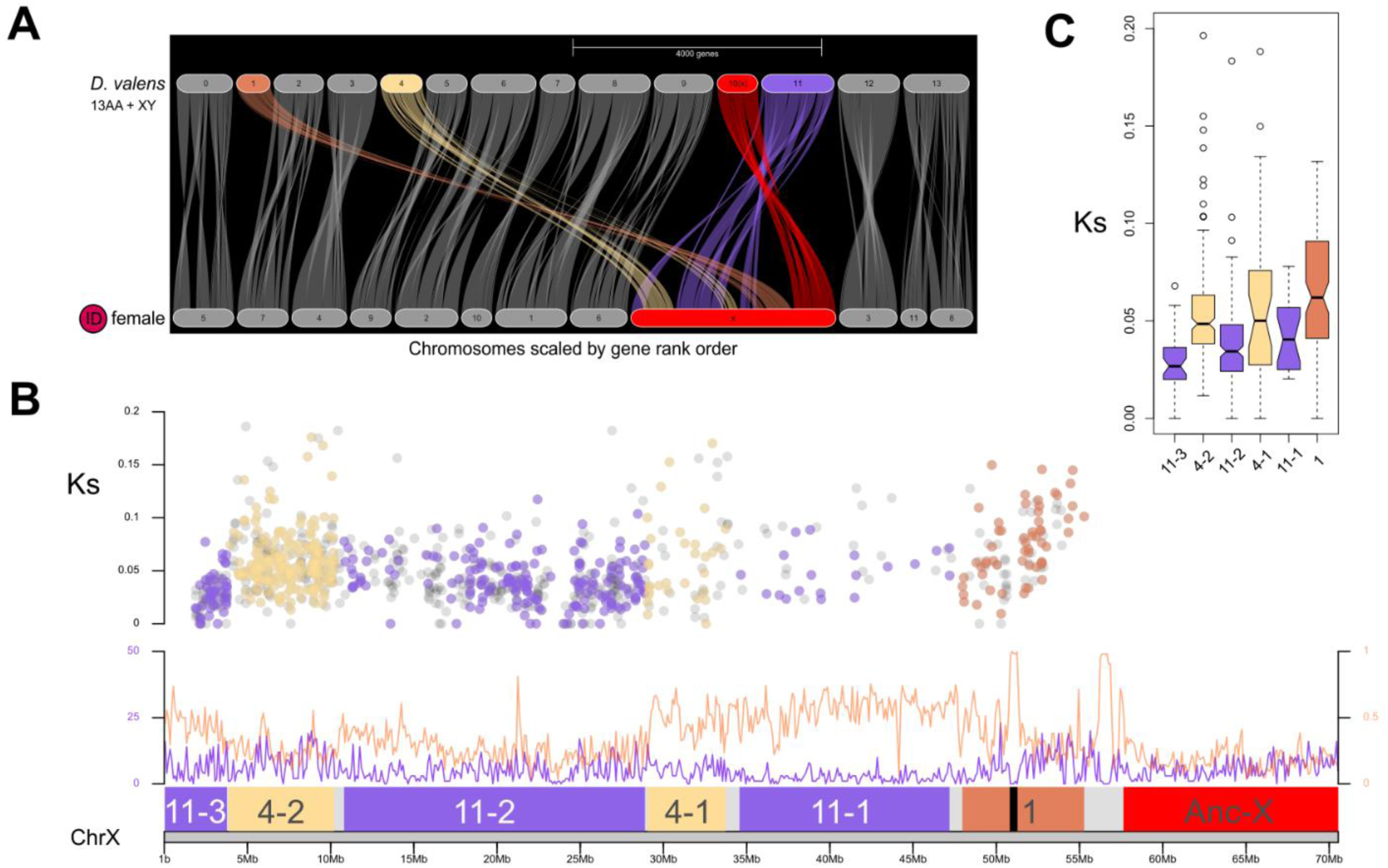
The neo-X in MPB formed via sequential fusions between the conserved X chromosome and ancestral autosomes. **A)** Synteny comparison between *Dendroctonus valens* and representative ID female MPB. X chromosomes shown in red. Autosomes in *D. valens* that are X-linked in MPB are highlighted. **B)** Synonymous substitution rate (Ks) for neo-X/neo-Y gametologs plotted using neo-X coordinates with points color-coded by *D. valens* autosomal location (top panel). Grey points are MPB gametologs without clear orthologs in *D. valens*. Neo-X chromosomal structure (bottom panel) showing repetitiveness and gene locations (similar to Fig. 1D) and putative centromere location (black bar). Each unique block of ancestral autosome was sequentially numbered from the Anc-X for subsequent analyses to disintengle the history of the neo-sex chromosomes. Outlier Ks estimates above 0.2 not shown. **C)** Boxplot of gametolog Ks estimates by block and colored by *D. valens* autosome. Outlier Ks estimates above 0.2 not shown.

Although several possible scenarios exist for how the large neo-X chromosome formed, the two most plausible and distinct are: 1) several autosomes fused in an ancestor forming a single chromosome, which then fused to the anc-X in a single event, with the homologue becoming the neo-Y, or 2) over time, through sequential addition, autosomes have fused to the neo-X (and neo-Y). One way to test these scenarios is to compare the synonymous substitution rate, Ks, between neo-X/neo-Y gametologs to estimate when recombination ceased and explore spatial overlap with our observed synteny patterns [41, 42]. A noted challenge here is that we are only able to spatially localize when recombination ceased, and this approach therefore does not provide an estimate of when specific autosomes fused. Estimates from 522 MPB gametologs with *D. valens* orthologs reveal a pattern more consistent with a series of sequential fusion events, interrupted by periods during which recombination was lost; the signal of which appears stronger for some fusions (**Fig. 3B**). We found Chr1 gametologs had the highest median Ks, and this ancestral autosome is immediately adjacent to the ancestral X, and further, appears to currently harbor the centromere. For Chr4, we found a lower median Ks for both ancestry blocks (Chr4-1, Chr4-2) that were nearly equivalent, suggesting this chromosome likely lost recombination while still a single syntenic block, and subsequent inversion(s) separated Chr4. Chr11 (1-3), has the lowest median Ks estimates with slight differences between ancestry blocks suggesting this ancestral autosome lost recombination more recently and possibly in some stepwise fashion (**sTable 6**). Using median Ks estimates from each ancestral autosome, we date the loss of recombination between neo-X and neo-Y fusions as ∼ 8.6 MYA for Chr1, ∼ 6.3 MYA for Chr4, and ∼ 4.3 MYA for Chr 11 (**sTable 6**). These estimates provide additional support for the hypothesis that the neo-sex chromosomes formed prior to the split from sister species *D. jeffreyi* [11]. These analyses also revealed that the putative centromere is in a region ancestrally autosomal (Chr1), and the X chromosome inversions that distinguish our OR, ID and NV (**Fig. 2D**) are located entirely within a low-recombining region of the neo-X chromosome [17].

### Neo-X vs neo-Y differentiation and degeneration

Given the estimated ages when recombination stopped between neo-X and neo-Y, we next sought to characterize the extent of structural divergence. We found that only ∼65% of the neo-Y could confidently be aligned to the neo-X across all three Y haplogroups (**Fig 4A, sFig. 8**). Further, we found a profusion of inverted sequences in regions that could confidently be aligned (**Fig 4A, sFig. 8**). Accurately quantifying and establishing the evolutionary history of such numerous and complex structural rearrangements is exceptionally challenging [43] and our preliminary exploration revealed commonly used software struggled to properly enumerate and classify the observed differences. Therefore, we took a simplified approach and counted inverted alignments between neo-X and neo-Y, accommodating for missing/deleted (and therefore unalignable) regions of the neo-Y. In these contrasts in ID, NV, and OR, we estimated 1,034, 958, and 1,240 inverted segments, respectively (**sTable 7**). Many inverted segments were small (∼56% 1kb-10 kb). Moderate and large inverted segments were less common (∼42% 10 kb-100 kb, ∼2% > 100 kb) (**Fig. 4A**). Thus, although the precise inversion and translocation history remains unclear, the neo-Y is highly rearranged with respect to the neo-X, with only small tracts of conserved collinear sequence.

**Figure 4.**
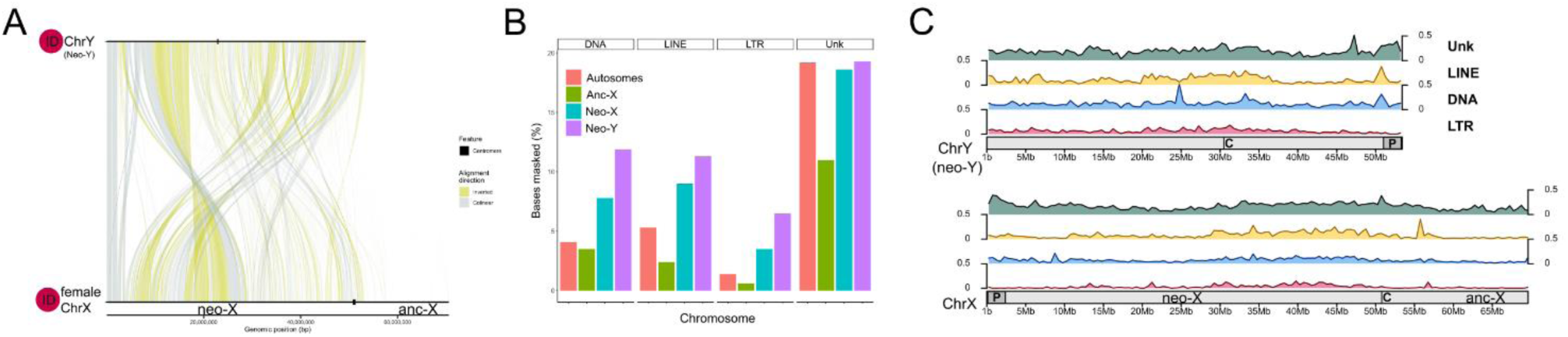
The neo-Y is highly differentiated structurally from the neo-X yet only moderately enriched for repetitive elements. **A)** Collinearity between ID neo-X and neo-Y. Putative centromere location shown with black bar and inverted sequence shown with yellow ribbons. Note the orientation of the neo-Y is flipped to align the PAR (far left). **B)** Percentage of bases masked for specific repeat families by chromosome type. **C)** The proportion of bases masked for repetitive sequence in 50 kb windows plotted across the entire X (ancestral and neo) and neo-Y. The most common families of repeats are shown (LINEs, DNA transposons, LTRs) as well as unknown repeats. The PAR (P) and centromere (C) locations are denoted.

Repetitive sequences, and specifically TEs, are expected to expand the size of young sex chromosomes, and specifically the neo-Y [5, 8, 44] and consistent with this, we found 49.3% of bases were repeat masked for neo-Y compared to 38.9% for the neo-X (**sTable 8**). Interestingly, the anc-X has far fewer TEs than the neo-X, with only 17.6% of bases masked for TEs. The autosomal average was found to be 30.5% but differed by autosome. Of the repeat families, LTR elements showed the most substantial difference when comparing the neo-X to neo-Y (3.5% vs 6.5%)(**Fig. 4B and C, sTable8**). Thus, although the assembled length of the neo-X and neo-Y are quite similar, we find an increased prevalence of TEs on the neo-Y.

We reasoned that the observed structural changes and repeat proliferation on the neo-Y likely resulted in Y-specific changes that complicated annotation and would hinder a full accounting of functional and non-functional genes. Therefore, we used Orthofinder [45] to identify gene relationships between our three male genome annotations and used Liftoff [46] to map neo-X genes back to the neo-Y. We focused our analyses on the ID male genome because of its contiguity and attempted to lift 2,898 annotated neo-X genes over to the neo-Y. We failed to lift ∼22% genes (*n* = 645), suggesting these were completely absent, while an additional ∼25% (*n* = 716) lifted over but failed to overlap an annotated neo-Y gene and/or had <50% sequence identity. In addition, ∼11% (*n* = 302) of genes lifted over but hit multiple neo-Y genes, many of which upon inspection appeared to be small gene fragments. We omitted ∼3% (*n* = 98) of genes as they were in complex relationships that did not allow us to assess degeneration. In total, we were able to confidently lift 39% of genes (*n* =1,137) over to a location where a single annotated neo-Y gene resided, which we therefore considered gametologs. Through multiple sequence alignments of these gametologs, we found that ∼11% (130 genes/1,137 alignments) of these neo-Y copies had frameshift mutations and/or internal stop codons. Thus, in total, we find that ∼62% of neo-Y genes show some form of degeneration, ranging from gene deletions to mutations disrupting reading frames.

### Identification of pseudoautosomal region (PAR) and evidence of ongoing degeneration

To maintain proper migration during meiosis, heteromorphic sex chromosomes often synapse in the heterogametic sex (males in XY taxa) in a small region of the chromosome known as the pseudoautosomal region (PAR). Interestingly, many beetle species with highly heteromorphic sex chromosomes pair the X and Y at a distance during meiosis and lack PAR(s) [47], including some *Dendroctonus* without neo-sex chromosomes [40]. To determine if MPB had putative PAR(s), we scrutinized the Hi-C contact maps from the three males to identify associations between sex chromosomes. We identified a small region on one end of the neo-X and neo-Y enriched for contacts as predicted for regions with underlying sequence similarity and/or are in close physical proximity (**sFig. 1, sTable9**). Male and female Illumina and PacBio HiFi sequencing coverage analysis of these regions showed clear convergence in coverage, in contrast to the rest of the sex chromosomes, consistent with this region being the PAR (**sFig. 3**). Four of the six assembled neo-X chromosomes contained PARs captured in long contigs (> 40 Mb) (**sFig. 2**). Additionally, all three neo-Y PARs were assembled as either one or two contigs that were scaffolded via Hi-C to the end of the neo-Y. The end of each independently assembled and scaffolded neo-Y PAR had terminal telomeric sequence, thus providing additional evidence of the correct orientation. Thus, we were able to confidently assemble and identify one PAR on each neo-X and neo-Y.

The PARs from OR and ID were found to be of similar length (∼2.17 Mb) and gene content (∼123 protein coding genes). In contrast, the NV PAR was found to be 8% shorter (∼1.99 Mb) and had 23 fewer protein coding genes (96 total). Genes missing from the NV neo-Y PAR were found in a single alternate location on the NV neo-Y, seemingly the result of past inversions (**sFig. 9**). Interestingly, all assembled X chromosomes were found to be largely collinear for the first ∼50 Mb of the chromosome which includes the PAR (**Fig 2A**, **sFig 9**), suggesting structural changes specific to the NV neo-Y have shortened the NV PAR. Consistent with this hypothesis, seven high-quality gametologs that appear to have moved outside of the PAR in NV were found to have more synonymous substitutions than those that still remain in the PAR in ID and OR (NV = 2.3% ± 0.6, ID = 0.9% ± 0.4, OR = 1.4% ± 0.5) suggesting ongoing degeneration and erosion of the PAR.

### Segregating Neo-Ys are structurally diverse and uniquely degenerated

To broadly characterize neo-Y sequence similarity between the three Ys, we first used whole chromosome alignments and were able to confidently align ∼91% of sequence in all pairwise comparisons. Despite this fairly high alignment, we found abundant structural variation and using methods outlined above, found 291 inverted alignments between OR and ID, 341 between NV and ID, and 434 between NV and OR (**sTable 10**). We classify more than half of these as moderate to large inverted segments (∼61% 10 kb-100 kb, ∼18% > 100 kb). While the exact number and sequence of events that led to this remarkable level of structural variation is difficult to reconstruct, this profusion hinted at broadscale changes that could impact gene function and content.

The extensive rearrangements made assessing neo-Y gene content exceptionally challenging, so we took a two-step approach to help identify genes and characterize pseudogenization. We first used OrthoFinder [45] to identify gene relationships using the predicted proteins from our three neo-Y annotations. Similar to Moraga et al. [48], we took a conservative approach and examined genes identified as found on all (1:1:1), missing from one (1:1:0), and missing from two (1:0:0) neo-Ys. In total, we were able to retain 2,082 genes (∼87%) with 51% (*n* = 1061) categorized as 1:1:1 based purely from our *de novo* annotations. We then looked for degenerated copies (pseudogenes) on missing neo-Ys using Liftoff [46]. We were able to promote 79% of genes (374/472) from missing from one (1:1:0) to found on all (1:1:1) and promoted ∼94% of genes (518/549) from missing from two (1:0:0) to missing from one (1:1:0, ∼15%, *n* = 84) or found on all (1:1:1, ∼79%, *n* = 434). This process allowed us to recover ∼90% of genes or pseudogenes (*n* = 1,869) on all three neo-Ys, while ∼9% (*n* = 182) were still found as missing from one neo-Y, and ∼1% (*n* = 31) were missing from two neo-Ys (**Fig. 5A**). For contrasts where we could assess sequence degeneration and pseudogenization between neo-Y copies (i.e., 1:1:1 and 1:1:0 categories), we estimated the proportion of genes with at least 1 gene copy containing a frameshift mutation and/or internal stop codon. We found that ∼10% of neo-Y genes had at least one copy that showed some evidence of degeneration (**Fig. 5B**). This was significantly higher than other chromosomes (**sTable 11**). As expected, frameshift mutations and internal stop codons were overrepresented in lifted neo-Y gene copies (∼28% of gene copies) compared to the *de novo* annotated neo-Y gene copies (≤0.5%). Our results suggest that gene lifting was a useful method for identifying degeneration and that these genes were likely missed during annotation because they lacked sufficient sequence similarity or transcriptomic evidence.

**Figure 5.**
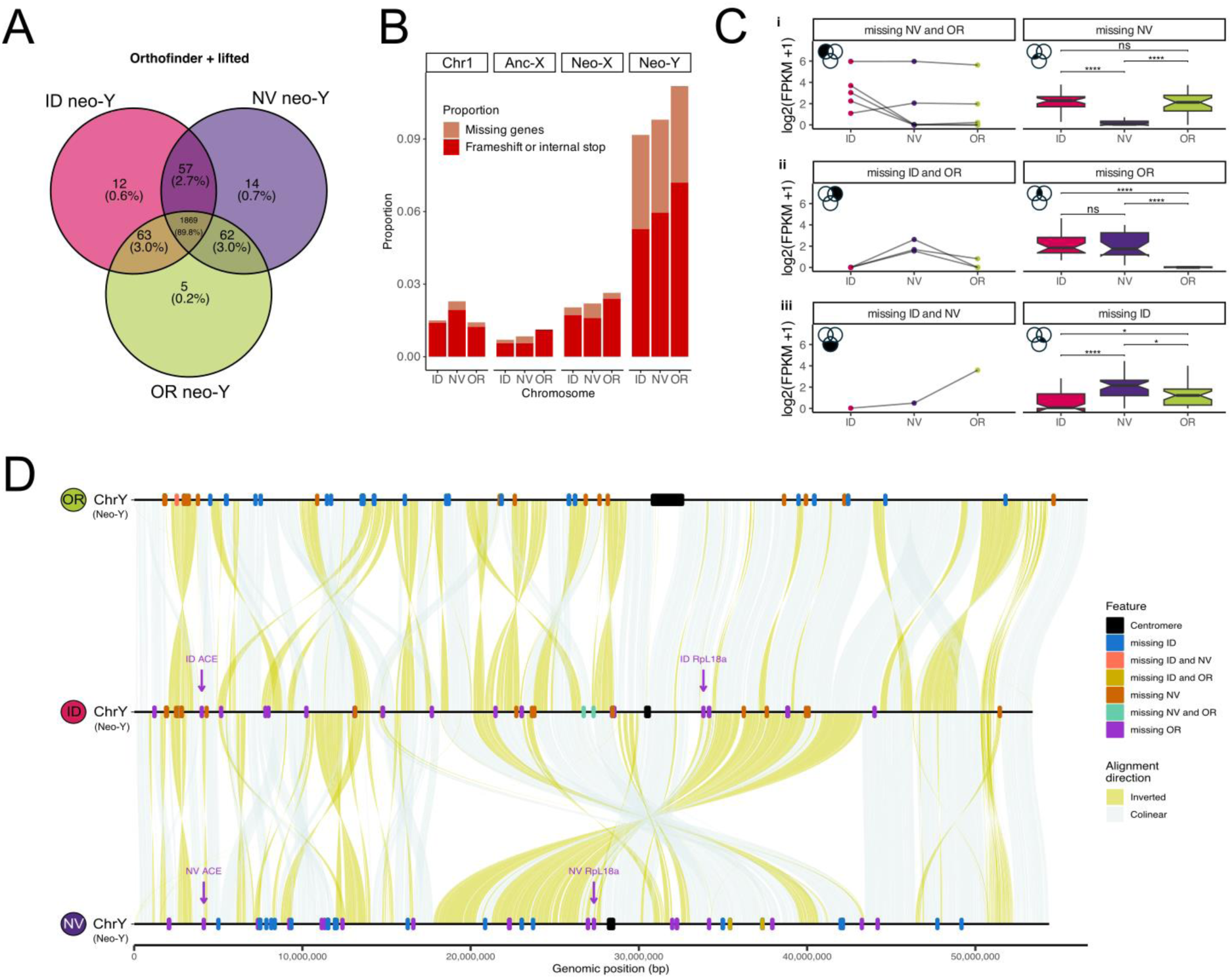
Neo-Ys differ in structure, gene content, and function. **A)** Overlap of 2,082 genes present across the three neo-Y haplogroups. **B)** Neo-Y degeneration was revealed by assessing missing genes and interrogating multiple sequence alignments. The neo-Y had a signficantly more missing genes and frameshifts or premature stops than the neo-X and anc-X (for gene counts and significance tests see **sTable 11**). **C)** Median expression from three testes replicates per neo-Y haplogroup. Each row represents expression relative to the three neo-Y assemblies (i = ID, ii = NV Y, iii = OR). Outliers were excluded from the plot (see **sTable 12** for signficance tests, ns = not significant (p > 0.05), ** = p ≤ 0.05, ** = p ≤ 0.01 *** = p ≤ 0.001 **** = p ≤ 0.0001) **D)** Collinearity between the three assembled neo-Ys. Missing genes with predicted functions are represented by color-coded ticks along the chromosome. Note the PAR is oriented to the right. See **sFig. 10** for plots reflecting autosomal ancestry.

We then turned our attention to the remaining 314 gene families (13%), which we had identified as multi-gene copies and gene fragments. We focused our search on neo-Y gene expansions using functionally annotated gene families (∼63%, *n* = 197) that were present with at least one copy on the neo-X and were multicopy on the neo-Y. In total, we found 17 possible gene expansions shared across all three neo-Ys, 7 were shared between two neo-Ys, and 2 were private (**sTable 13**). Of these gene expansions, only forkhead box F1 protein (*FOXF1*) was a gene expansion of ≥ 5 gene copies across all three neo-Ys. *FOXF1* expanded from a single neo-X copy to 8-13 Y copies (**sTable 12**), likely via tandem duplications based on proximity to each other (**sFig. 12)**. There was a lack of widespread gene proliferation on the neo-Y and we failed to find clear signals of gene expansions as seen in other neo-sex chromosome systems [49, 50]

### Functional relevance of missing Y genes

To confirm missing neo-Y genes were not due to genome assembly issues, and that retained neo-Y genes were still transcriptionally active, we re-examined our male and female transcriptomic data. By mapping reads from both sexes to each male genome assembly with our lift-over aided annotations, we could restrict analyses to neo-Y genes (or sections thereof) with sufficiently divergent sequence from their neo-X gametolog, thereby allowing us to characterize neo-Y gene expression. We were able to assess neo-Y gene expression for 61, 60, and 64 genes for ID, NV, and OR, respectively (**Fig 5C**); ∼72% of which were functionally annotated. Genes that were identified as either missing from NV or missing from OR showed almost no evidence of expression (**Fig. 5Ci and 5Cii**) suggesting these genes are indeed absent. Genes identified as missing from ID had significantly lower expression than the same gene in NV and OR (**Fig. 5Ciii**). Most missing ID genes had no expression (*n* = 22), while we detected some expression from 12 genes. Expression was generally lower in genes that were missing from two neo-Y haplogroups (e.g., missing NV *and* OR) **(Fig 5Ci).** Investigation of missing genes with atypical expression patterns showed that reads were not mapping to all exons and/or the genes were adjacent structural variation **(Fig 5D)** suggesting these missing genes were likely gene fragments. These analyses were robust to reference genome choice (**sTable12**) illustrating that the genes classified as missing are indeed absent.

Previous research identified several highly expressed neo-Y genes that were present in Eastern and Central, yet missing from Western, and therefore possibly play a role in the observed hybrid sterility in MPB [11]. The most promising candidates were angiotensin-converting enzyme (*ACE*) and 60S ribosomal protein L18a (*RpL18a*). We were able to confirm that *ACE* and *RpL18a* are indeed missing from the OR Y and yet present and expressed in ID and NV (**Fig. 5D, sFig. 13).** *RpL18a* is a haploinsufficient gene in *Drosophila melanogaster* and low dosage leads to developmental issues [51, 52] while *ACE* genes have been found to play a role in spermatogenesis [53]. Interestingly, *ACE* and *RpL18a* became X-linked with the oldest fusion (Chr1) **(Fig. 3B)**, and further, the neo-X copies are in the population-specific inversions bordering the neo-X centromere **(Fig. 2D)**. In total, our analysis identifies 27 genes completely absent from the OR neo-Y. Further, we identified a combined 77 new candidate genes across all three neo-Ys with reported functions that will serve as promising candidates for studying reproductive incompatibilities and neo-X/neo-Y gene expression regulation between populations (**Fig. 5D**). Many of these genes are adjacent structural differences between neo-Ys suggesting inversions and deletions are modifying the gene landscape with numerous transcriptionally active genes missing from specific neo-Y chromosomes.

## CONCLUSIONS

Our results provide a new window into the early stages of neo–sex chromosome formation and neo-Y chromosome degeneration (**Fig. 6A**). Importantly, our study demonstrates that multiple neo-Y chromosomes, each with numerous and distinct structural and likely functional differences, can coexist within a single species (**Fig. 6B**). The sex chromosome–autosome fusions that we estimate occurred millions of years ago appear to have set the stage for extensive neo-Y degeneration, ultimately producing three distinct neo-Ys in MPB. Prior to the degeneration characterized here, we estimate that the ancestral Y chromosome contained more than 2,000 functional genes that were ancestrally autosomal in other closely related *Dendroctonus* species.

**Figure 6.**
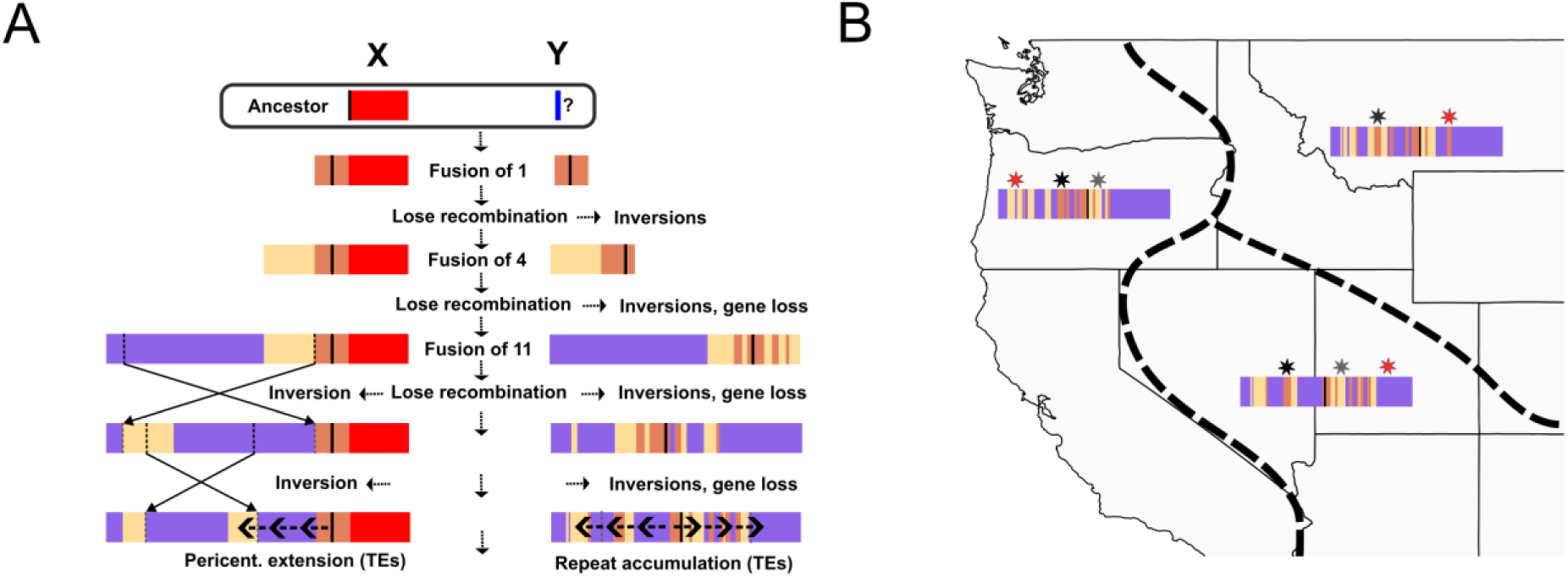
Evolutionary model of MPB sex chromosomes. **A)** Our results suggest the following events led to the present day neo-sex chromosomes in MPB. In the ancestor of MPB (and sister species *D. jeffreyi*), ancestral autosome 1 fused to the X and its homologue became the neo-Y, possibly via fusion with an ancestral Y. Inversions likely initiated the loss of recombination between the neo-X and neo-Y, and subsequent inversions proliferated on the neo-Y. Autosome 4 then fused, and followed a similar trajectory of recombination loss, and inversion proliferation. Autosome 11 then fused. On the neo-X, two inversions seperated 4 and then 11. The region adjacent the current centromere (located in ancestral autosome 1) has since experienced expansion of transposable elements (see Fig. 2) common in pericentromeric regions. For the neo-Y, once recombination ceased with any neo-X, inversions appear to have proliferated at a rate far exceeding the neo-X, and gene loss ensued. Transposable elements are just beginning to expand along the chromosome and have yet to reach densities typical of evolutionarily old Y chromosomes. **B)** Ongoing degeneration of the neo-Y chromosome is occurring in MPB populations. Each neo-Y type has numerous unique inversions and distinct gene loss. Degeneration is the result of mutations shared by all (black star), shared by two (grey star) or unique to a neo-Y (red star).

Despite these striking patterns, little is known about natural variation in neo-Y (or neo-W) chromosomes, particularly when large, gene-rich chromosomes are involved, or about how the rapid accumulation of mutations and pseudogenes affects fitness. Similarly, it remains unclear whether and how dosage compensation mechanisms evolve in response to these gene losses. Many questions therefore remain in this system. For example, does dosage compensation occur to buffer degenerated neo-Y genes, and if so, does it operate on a gene-by-gene basis or through a chromosome-wide mechanism [21]? Could mismatches between neo-X and neo-Y chromosomes lead to gene misregulation underlying the observed reproductive incompatibilities in MPB [10]? Intriguingly, recent models propose such mismatches as a potential link between sex chromosome evolution and well-known patterns in the genetics of speciation [9, 54]. Further investigation of gene regulation in MPB and in interpopulation hybrids will be particularly illuminating.

## MATERIALS AND METHODS

### Sample collection

Beetles were collected by felling infested trees at three separate locations in the summer of 2021. Tree sections were then transported to the USDA Forest Service Rocky Mountain Research Station in Logan, UT, and adult beetles were reared out of the host material. The western population was collected from an infested lodgepole pine (*Pinus contorta*) near Sumpter, OR (44°44’43.85”N, 118°23’46.42”W). The eastern population was collected from lodgepole pine (*Pinus contorta*) near Fairfield, ID (43°32’35.27”N, 114°49’7.63”W). The central population was collected from limber pine (*Pinus flexilis*) near Jarbidge, NV (41°49’42.01”N, 115°28’4.11”W). Adult beetles were sexed using morphological features [55].

### Long-read genome sequencing

Flash frozen and sexed adult beetles were sent to the USDA-ARS Tropical Pest Genetics and Molecular Biology Research Unit in Hilo, HI. High molecular weight DNA was extracted from individual beetles using a Qiagen MagAttract HMW DNA Kit (Qiagen, Hilden, Germany). The Genomic DNA 165kb Kit on an Agilent Femto Pulse system and Qubit Broad Range dsDNA kit on a Denovix DS-11 along with UV spectrophotometric readings were used to qualified and quantified the DNA. The resulting DNA from a single beetle was subjected to shearing on a Diagenode Megaruptor 3, targeting 15 - 20 kb fragments and prepared into a barcoded PacBio HiFi sequencing library using a SMRTBell Express Template Prep 3.0 kit. The resulting library was sequenced on a partial SMRT cell of PacBio Revio system.

### Hi-C library construction and sequencing

For Hi-C sequencing, a separate individual from the same sex and same population matching the HiFi long-read sample was also flash frozen in liquid nitrogen and stored at -80C before preparation and sequencing. The Hi-C library was prepared following a standard protocol using the restriction enzymes DdeI and DpnII, and the resulting proximity ligation library was prepared for sequencing using the NEBNext Ultra II DNA Library Prep Kit. The final libraries were sequenced on a partial flow cell using the AVITI 2×150 Sequencing Kit Cloudbreak FS High Output kit on the Element AVITI System. Following sequencing, raw reads were basecalled using bases2fastq v.1.3.0.

### De novo genome assembly and Hi-C scaffolding

Raw HiFi reads were processed prior to assembly using HiFiAdapterFilt v2.0 [56]. The resulting filtered reads were then assembled into primary and alternate assemblies using HiFiASM and the --primary setting [57]. The contigs from the primary assembly were used for all autosomes, while both the primary and accessory assemblies were interrogated for sex chromosome contigs since undifferentiated regions, like the PAR, would be present in both. Hi-C reads were mapped to the genomes using BWA MEM [58] with all intermediate files being modified using samtools version1.17 [59] with duplicates removed via Picard version 3.0.0 MarkDuplicates (http://broadinstitute.github.io/picard). YaHS [60] and Juicer [61] were used to create Hi-C contact maps to assist with scaffolding the genome. Output files were visualized, and the genome assembly was manually curated and putative chromosomes ordered using Juicebox and associated scripts [62]. Genome completeness was determined using BUSCO [63] and the endopterygota odb10 database. In male assemblies, over ∼11% of BUSCOs were identified as “duplicated”, and as expected, nearly all of which were later identified as gametologs that reside on both the neo-X and neo-Y (**sTable 2**).

### Repeat identification

To identify repetitive elements we used our *de novo* genome assemblies as the input database in repeatmodeler2 version 2.0.5 [64]. The resulting consensus repeat families were then used to mask their respective genome using RepeatMasker version 4.1.5 and the -no_is -nolow -xsmall flags. We used TRASH [37] to identify consensus satellite sequences and plot their relative locations in each reference genome. To further evaluate the candidate centromere-associated satellite sequences, we used self-identity comparisons in ModDotPlot [38], with default settings to visualize repetitive patterns across the chromosomes. We used KaryoploteR [65] to visualize the location of the core consensus sequence in the OR male genome.

### Genome annotation

To aid annotation, we generated transcriptomic data from several beetle tissues, including male/female heads, ovaries, and testes, from the OR, ID, and NV populations. Beetle tissues were isolated from live adults in phosphate buffered saline and quickly transferred to TRIzol (Invitrogen) for stabilization. We then used the Direct-zol RNA MiniPrep (Zymo Research) kit and standard methods to extract total RNA. RNA quality and quantity were assessed via a High Sensitivity Screen on an Agilent 2200 TapeStation prior to sequencing. RNAseq libraries were prepared at the Indiana University Center for Genomics and Bioinformatics using an Illumina TruSeq Stranded mRNA kit. The resulting libraries were sequenced on an Illumina NextSeq 2000 for 200 cycles.

To create the input files required for the BRAKER3 pipeline, raw Illumina RNAseq reads were first processed using SeqyClean version 1.10.09 using recommended settings with poly-A/T trimming enabled and minimum read length 30bp [66]. Trimmed reads were then mapped to their respective reference genome using hisat2 version 2.2.1 [67] with default settings, then converted to bam files and indexed using samtools version 1.17 [59]. We then used BRAKER3 [28, 68, 69] to independently annotate the six soft-masked primary genome assemblies. To help train the software to identify protein coding genes, we used predicted proteins from the published *Dendroctonus ponderosae* assembly and annotation (NCBI RefSeq assembly GCF_020466585.1; [19]) and the appropriate gene expression data for each lineage/sex combination outlined above.

### Synteny and collinearity comparisons

To explore broad patterns of synteny and collinearity between our six beetle genome assemblies, we used the predicted proteins from our initial annotations and Genespace [30] with default settings. For additional fine-scale, chromosome alignment comparisons, we used minimap2 version 2.26 [30, 31] with the following parameters: -x asm20 -c --eqx --secondary=no. We estimated mapping rates between query and references by using the length of aligned query sequence and dividing it by the reference chromosome size. We visualized alignments with SVbyEye version 0.99 [70].

To estimate the number of structural rearrangements between neo-X and neo-Y, we counted the number of inverted alignments > 1,000 bp when comparing the neo-Y to its corresponding female neo-X. We merged neighboring neo-Y inverted segments when the gap between neighbors was < 25 kb on the neo-X. We considered this a conservative approach that was done to account for missing/deleted regions on the neo-Y that interrupted regions that were still mostly collinear. We estimated structural rearrangements between neo-Ys by using an OR reference vs an ID query, an ID reference vs an NV query, and an OR reference vs an NV query. Reference and query genomes were swapped and produced similar results.

### Exploring evolutionary strata

We first used Genespace [30] to compare the ID female genome assembly to *Dendroctonus valens*, a more distantly related species without neo-sex chromosomes, and a karyotypic formula (13AA + Xyp). Phylogenetic analyses suggest this species is likely a good representative of the ancestral chromosomal configuration in *Dendroctonus* [71]. For *D. valens*, we used a recently published chromosome-level assembly and annotation [39]. To estimate the synonymous substitution rate, Ks, for neo-X and neo-Y gametologs, we used the results from Orthofinder (above) to identify 1:1 gametologs between the ID neo-X and neo-Y. We then used PRANK [72] to perform codon-aware alignments of the CDS for each gametolog, and KaKs_Calculator [73] to estimate Ks. The number of gametologs, their relative genomic location in *D. valens*, and per gene Ks estimates are in (**sTable 6**).

To estimate putative dates of loss of recombination between chromosomes that fused to the neo-X and neo-Y, we used methods outlined in Krasovec et al [42]. Briefly, we used the equation t_g_ = Ks/2µ, where t_g_ is the time of divergence in generations, Ks is synonymous sequence divergence between gametologs, and µ is the per generation point mutation rate. We calculated the median Ks estimates for each ancestral autosomal or combined blocks (e.g., Chr4-1, Chr4-2) and report the per chromosome average (e.g., Chr4) (see **sTable6**). Since there is no estimated spontaneous mutation rate in beetles (order Coleoptera), we used an average rate estimated from three other holometabolous insect orders, yielding a rate of 3.93 x 10^-9^ (Diptera = 5.49 x 10^-9^ [74], Lepidoptera = 2.90 x 10^-9^ [75] Hymenoptera = 3.40 x 10^-9^ [76]). *Dendroctonus ponderosae* typically has one generation a year [77].

### Identifying degenerated neo-Y genes

We first used Orthofinder [45] and the longest predicted protein from each gene from the three *de novo* annotated Y chromosomes, Y genes were then binned into one of four categories: “core” (1:1:1, on all three Y), “shared” (1:1:0, missing from a single Y), “private” (1:0:0 missing from two Ys), or “multi” (complex multi-copy families due to duplication and fragmentation). We also performed the same categorization with Orthofinder for Chr1 and the neo-X. These chromosomes had high levels of gene synteny (**Fig 2A**) and chromosome collinearity (**sFig 5**) so we would expect genes to be consistently identified between them.

After noticing a discrepancy in proportion of shared and private genes on the neo-Y compared to Chr1 and ChrX (**sFig. 11**), we wanted to ensure that the *de novo* neo-Y annotations were not missing genes that were present at the nucleotide level but not expressed in the sequenced tissues. We used Liftoff version 1.6.3 [46] to lift “shared” and “private” genes with default parameters and specified -s 0.70, -exclude_partial and -polish. We specified ≥70% sequence identity compared to the default ≥50% sequence identity threshold so we were more likely to lift genes that were functionally present on a degenerating neo-Y. We used these lifted annotations to further scrutinize the categories described above. We promoted shared genes to a core neo-Y gene if at least one copy of the shared gene was lifted to the missing neo-Y. Private genes that were lifted were promoted to shared or core neo-Y genes. For comparison, we followed the same promotion methods for “private” and “shared” genes on Chr1 and the neo-X. We refer to these male annotations as the “lift-over aided annotation” throughout the manuscript. The remaining complex multi-copy gene families were excluded from Liftoff evaluation (Chr1 = 74, X = 235, Y = 314). We visualized gene count results using ggvenn version 0.10.0 [78].

### Characterizing degeneration between the neo-X and neo-Y

To identify neo-X/neo-Y gametologs, we mapped neo-X genes to the neo-Y. We used Liftoff version 1.6.3 [46] with default parameters but specified ≥70% sequence identity (-s 0.70). We chose Liftoff because it uses minimap2 [30, 31] which takes into account gene structure to facilitate mapping to complex regions. Neo-Y genes were considered “absent” if we failed to lift over the neo-X gene. Next, we intersected the lifted neo-X gene coordinates against the lift-over aided neo-Y annotation in BEDtools version 2.31.0 [79]. We assessed degeneration using the following criteria, 1) neo-X genes that were lifted onto the neo-Y but failed to meet the default mapping coverage (≥50%) or sequence identity (≥70%) (potential pseudogenization), 2) neo-X genes that did not overlap with any coordinates of an annotated neo-Y gene (gene loss events), and 3) intersected with multiple neo-Y genes (potential gene fragmentation). Neo-X genes that overlapped with a single neo-Y gene were retained for further assessment of sequence changes. Neo-X genes that did not fit any of these criteria were omitted from further analyses. We examined gametologs for frameshifts or internal stop codons by aligning their coding sequences in MASCE version 2.07 [80]. We considered the neo-X genes reliable (--seq) and the neo-Y genes as less reliable (--seq_lr) since neo-Y genes were likely to align poorly due to degeneration. We also increased the cost of frameshifts for reliable sequences (--fs 80 and --fs_term 80) since the neo-X genes were likely to remain intact compared to their gametologs. We evaluated the sequences with the output generated from “-out_stat_per_seq”.

### Assessing gene loss and sequence degeneration between the three neo-Ys

To assess degeneration between the respective neo-Ys, we assessed gene loss and sequence degeneration. We also performed the same analyses for Chr1, neo-X, and anc-X as a point of comparison. We quantified the number of uniquely missing genes – these are genes that are absent from a neo-Y haplogroup because the genes are either private or shared. We accounted for uniquely missing genes on Chr1 and ChrX that may have resulted from regions that were not assembled by evaluating the presence of the homologous region across the three assemblies by omitting them from the proportions.

To assess sequence degeneration, we quantified frameshift mutations and internal stop codons using the shared and core genes. We used the predicted coding sequences from our liftover-aided annotation but needed to extract the lifted genes for the final list of coding sequences. We extracted the coding sequences from lifted genes with the “agat_sp_extract_sequences.pl” script from AGAT version 1.4.1 [81]. We aligned sequences for shared and core genes with MASCE version 2.07 [80] and specified the following flags for frameshift penalties (--fs 80 and --fs_term 80). We quantified the proportion of genes with internal frameshifts or internal stop codons for the neo-Y. We assessed statistical significance between chromosome types using a test of equal proportions with the “prop.test” function in R version 4.5.0 [82].

### Evaluating gene expansions on sex chromosomes

To understand the functional relevance of MPB genes, we functionally annotated all our lift-over aided annotations using the proteins from *Dendroctonus ponderosae* (GCF_020466585.1, [19]) and *Tribolium castaneum* (icTriCast1.1; GCF_031307605.1). We built a DIAMOND database [83] using these protein sequences and used the file as input for eggNOG emapper-2.1.12 with default parameters [84]. To simplify gene expansion analyses, we focused on gene families that were functionally annotated. If at least one gene from the gene families was functionally annotated, the gene family was labeled as a functionally relevant and retained for downstream analyses. We excluded gene families that were not functionally annotated or had uncharacterized functions. We enumerated gene copy number for each functionally relevant gene families on the neo-X and neo-Y. We considered gene expansions when genes were present with at least one copy on neo-X and were multicopy on the neo-Y. We examined notable gene expansions in the self-identity comparisons generated in ModDotPlot [38].

### Validating missing neo-Y genes

To evaluate putatively missing neo-Y genes, we mapped tissue-specific RNA-seq reads from female heads and ovaries and male head and testes to each male genome assembly using hisat2 version 2.2.1 [67]. We sorted and indexed the mapped reads with samtools version 1.17 [59]. Using samtools view, we first attempted to restrict cross-mapping by excluding unmapped, secondary, and supplementary alignments (-F 2308) and retained properly paired reads (-f 2) with a mapping quality ≥30 (-q 30). We used StringTie version 2.2.3 [85] to evaluate FPKMs per gene with the final lift-over aided neo-Y annotations for each genome. We omitted any neo-Y genes with female FPKM ≥1 as these could not be assessed confidently due to cross mapping between the gametologs. For each neo-Y population, we calculated the median FPKM per missing gene and retained genes with a median FPKM ≥ 1 in at least one neo-Y haplogroup. Genes below this threshold were considered to not be expressed in these tissues. These filters left us with ∼47% of missing genes for evaluation. We tested for significant differences in read counts between the neo-Y haplogroups using a pairwise Wilcoxon test and adjusted for multiple comparisons with a Bonferroni correction in R version 4.5.0 [82].

We searched for the genes *ACE* and *RpL18a* in our missing genes dataset since they had been previously identified as missing from OR males [11]. Both genes were identified as missing in OR prior to filtering genes that had any females with FPKM ≥ 1. However, *RpL18a* had minor levels of cross mapping in the ID and NV females for the neo-Y copy with OR females having even lower levels of cross mapping. Despite this cross mapping, OR male expression was still near absent (**sFig. 13**), thus we chose to retain *RpL18a* as a promising candidate gene

## ACKNOWLEDGEMENTS

We thank the Indiana University Center for Genomics and Bioinformatics for assistance with RNA-seq library preparation and sequencing. Additional genome sequencing resources were supported by USDA’s Agricultural Research Service project numbers 2040-30400-003-000D, and USDA computation support through 0201-88888-003-000D, and 0201-88888-002-000D. This work would have been impossible without help from the Indiana University UITS high-performance computing cluster. This research was supported by NIH NIGMS MIRA grant: R35GM151123. All opinions expressed in this paper are the authors’ and do not necessarily reflect the policies and views of USDA. Mention of trade names or commercial products in this publication is solely for the purpose of providing specific information and does not imply recommendation or endorsement by the U.S. Government. USDA is an equal opportunity provider and employer. The authors declare no conflict of interest.

## DATA AVAILABILITY

Raw sequence reads and genomes are available under BioProject ID PRJNA1405530 via NCBI. Genome annotations, relevant code, and data are available at Figshare (https://doi.org/10.6084/m9.figshare.31073665).

## Notes

### Competing Interest Statement

The authors have declared no competing interest.

## REFERENCES

1. Charlesworth, B., The evolution of chromosomal sex determination and dosage compensation. Current Biology, 1996. 6(2): p. 149–162.

2. Bachtrog, D., A dynamic view of sex chromosome evolution. Current Opinion in Genetics & Development, 2006. 16(6): p. 578–585.

3. Rice, W.R., Evolution of the Y Sex Chromosome in Animals: Y chromosomes evolve through the degeneration of autosomes. BioScience, 1996. 46(5): p. 331–343.

4. Bachtrog, D., Y-chromosome evolution: emerging insights into processes of Y-chromosome degeneration. Nat Rev Genet, 2013. 14(2): p. 113–24.

5. Mahajan, S., et al., De novo assembly of a young Drosophila Y chromosome using single-molecule sequencing and chromatin conformation capture. PLOS Biology, 2018. 16(7): p. e2006348.

6. Zhu, Z., L. Younas, and Q. Zhou, Evolution and regulation of animal sex chromosomes. Nature Reviews Genetics, 2025. 26(1): p. 59–74.

7. Wei, K.H.C. and D. Bachtrog, Ancestral male recombination in Drosophila albomicans produced geographically restricted neo-Y chromosome haplotypes varying in age and onset of decay. PLOS Genetics, 2019. 15(11): p. e1008502.

8. Sacchi, B., et al., Phased assembly of neo-sex chromosomes reveals extensive Y degeneration and rapid genome evolution in Rumex hastatulus. Mol Biol Evol, 2024. 41(4).

9. Lenormand, T. and D. Roze, A single theory for the evolution of sex chromosomes and the two rules of speciation. Science, 2025. 389(6756): p. eado9032.

10. van der Bijl, W. and J.E. Mank, Hiding in plain sight: the Y chromosome and its reinvigorated role in evolutionary processes. Evolution Letters, 2024. 9(2): p. 165–171.

11. Bracewell, R.R., et al., Rapid neo-sex chromosome evolution and incipient speciation in a major forest pest. Nature Communications, 2017. 8(1): p. 1593.

12. Dowle, E.J., et al., Reproductive isolation and environmental adaptation shape the phylogeography of mountain pine beetle (Dendroctonus ponderosae). Molecular Ecology, 2017. 26(21): p. 6071–6084.

13. Bracewell, R.R., et al., Cryptic postzygotic isolation in an eruptive species of bark beetle (Dendroctonus ponderosae). Evolution, 2011. 65(4): p. 961–975.

14. Haldane, J.B.S., Sex ratio and unisexual sterility in hybrid animals. Journal of Genetics, 1922. 12(2): p. 101–109.

15. Presgraves, D.C., Sex chromosomes and speciation in Drosophila. Trends Genet, 2008. 24(7): p. 336–43.

16. Bentz, B.J., et al., Genetic architecture and phenotypic plasticity of thermally-regulated traits in an eruptive species, Dendroctonus ponderosae. Evolutionary Ecology, 2011. 25(6): p. 1269–1288.

17. Pushman, C., et al., QTL identification and characterization of the recombination landscape of the mountain pine beetle (Dendroctonus ponderosae). G3 Genes|Genomes|Genetics, 2025. 15(7).

18. Keeling, C.I., et al., Draft genome of the mountain pine beetle, Dendroctonus ponderosae Hopkins, a major forest pest. Genome Biol, 2013. 14(3): p. R27.

19. Keeling, C.I., et al., Chromosome-level genome assembly reveals genomic architecture of northern range expansion in the mountain pine beetle, Dendroctonus ponderosae Hopkins (Coleoptera: Curculionidae). Mol Ecol Resour, 2022. 22(3): p. 1149–1167.

20. Geneva, A.J., et al., Chromosome-scale genome assembly of the brown anole (Anolis sagrei), an emerging model species. Communications Biology, 2022. 5(1): p. 1126.

21. Bracewell, R., et al., Sex and neo-sex chromosome evolution in beetles. PLOS Genetics, 2024. 20(11): p. e1011477.

22. Xu, L., et al., Evolution and expression patterns of the neo-sex chromosomes of the crested ibis. Nature Communications, 2024. 15(1): p. 1670.

23. Peichel, C.L., et al., Assembly of the threespine stickleback Y chromosome reveals convergent signatures of sex chromosome evolution. Genome Biology, 2020. 21(1): p. 177.

24. Li, M., et al., Reconstruction of the origin of a neo-Y sex chromosome and its evolution in the spotted knifejaw, Oplegnathus punctatus. Molecular Biology and Evolution, 2021. 38(6): p. 2615–2626.

25. Muirhead, C.A., et al., Genomic origins and evolution of neo-sex chromosomes in Pacific Island birds. Proceedings of the National Academy of Sciences, 2025. 122(31): p. e2503746122.

26. Lanier, G.N. and D.L. Wood, Controlled mating, karyology, morphology, and sex-ratio in the Dendroctonus ponderosae complex. Annals of The Entomological Society of America, 1968. 61: p. 517–526.

27. Charlesworth, B., P. Sniegowski, and W. Stephan, The evolutionary dynamics of repetitive DNA in eukaryotes. Nature, 1994. 371(6494): p. 215–220.

28. Gabriel, L., et al., BRAKER3: Fully automated genome annotation using RNA-seq and protein evidence with GeneMark-ETP, AUGUSTUS, and TSEBRA. Genome Res, 2024. 34(5): p. 769–777.

29. Keeling, C.I., et al., Draft genome of the mountain pine beetle, Dendroctonus ponderosae Hopkins, a major forest pest. Genome Biology, 2013. 14(3): p. R27.

30. Lovell, J.T., et al., GENESPACE tracks regions of interest and gene copy number variation across multiple genomes. eLife, 2022. 11: p. e78526.

31. Dobzhansky, T. and A.H. Sturtevant, Inversions in the chromosomes of Drosophila pseudoobscura. Genetics, 1938. 23(1): p. 28–64.

32. Kapun, M., et al., Genomic evidence for adaptive inversion clines in Drosophila melanogaster. Molecular Biology and Evolution, 2016. 33(5): p. 1317–1336.

33. Noor, M.A.F., et al., Chromosomal inversions and the reproductive isolation of species. Proceedings of the National Academy of Sciences, 2001. 98(21): p. 12084–12088.

34. Gržan, T., et al., CenH3 distribution reveals extended centromeres in the model beetle Tribolium castaneum. PLoS Genet, 2020. 16(10): p. e1009115.

35. Veseljak, D., et al., Dynamic evolution of satellite DNAs drastically differentiates the genomes of Tribolium sibling species. Genome Res, 2025. 35(11): p. 2445–2460.

36. Prusakova, D., et al., Telomeric DNA sequences in beetle taxa vary with species richness. Sci Rep, 2021. 11(1): p. 13319.

37. Wlodzimierz, P., M. Hong, and I.R. Henderson, TRASH: Tandem Repeat Annotation and Structural Hierarchy. Bioinformatics, 2023. 39(5).

38. Sweeten, A.P., M.C. Schatz, and A.M. Phillippy, ModDotPlot—rapid and interactive visualization of tandem repeats. Bioinformatics, 2024. 40(8).

39. Liu, Z., et al., Chromosome-level genome assembly and population genomic analyses provide insights into adaptive evolution of the red turpentine beetle, Dendroctonus valens. BMC Biology, 2022. 20(1): p. 190.

40. Zúñiga, G., et al., Karyology, geographic distribution, and origin of the genus Dendroctonus Erichson (Coleoptera: Scolytidae). Annals of the Entomological Society of America, 2002. 95(3): p. 267–275.

41. Bergero, R., et al., Evolutionary strata on the X chromosomes of the dioecious plant Silene latifolia: Evidence From New Sex-Linked Genes. Genetics, 2007. 175(4): p. 1945–1954.

42. Krasovec, M., et al., The mutation rate and the age of the sex chromosomes in Silene latifolia. Current Biology, 2018. 28(11): p. 1832–1838.e4.

43. Alekseyev, M.A. and P.A. Pevzner, Breakpoint graphs and ancestral genome reconstructions. Genome Res, 2009. 19(5): p. 943–57.

44. Akagi, T., et al., Rapid and dynamic evolution of a giant Y chromosome in Silene latifolia. Science, 2025. 387(6734): p. 637–643.

45. Emms, D.M. and S. Kelly, OrthoFinder: phylogenetic orthology inference for comparative genomics. Genome Biol, 2019. 20(1): p. 238.

46. Shumate, A. and S.L. Salzberg, Liftoff: accurate mapping of gene annotations. Bioinformatics, 2021. 37(12): p. 1639–1643.

47. Blackmon, H. and J.P. Demuth, Estimating tempo and mode of Y chromosome turnover: explaining Y chromosome loss with the fragile Y hypothesis. Genetics, 2014. 197(2): p. 561–572.

48. Moraga, C., et al., The Silene latifolia genome and its giant Y chromosome. Science, 2025. 387(6734): p. 630–636.

49. Ellison, C. and D. Bachtrog, Recurrent gene co-amplification on Drosophila X and Y chromosomes. PLOS Genetics, 2019. 15(7): p. e1008251.

50. Bachtrog, D., S. Mahajan, and R. Bracewell, Massive gene amplification on a recently formed Drosophila Y chromosome. Nat Ecol Evol, 2019. 3(11): p. 1587–1597.

51. Marygold, S.J., et al., The ribosomal protein genes and Minute loci of Drosophila melanogaster. Genome Biol, 2007. 8(10): p. R216.

52. Cook, R.K., et al., The generation of chromosomal deletions to provide extensive coverage and subdivision of the Drosophila melanogaster genome. Genome Biol, 2012. 13(3): p. R21.

53. Hurst, D., et al., The Drosophila angiotensin-converting enzyme homologue Ance is required for spermiogenesis. Dev Biol, 2003. 254(2): p. 238–47.

54. Filatov, D.A., The two “rules of speciation” in species with young sex chromosomes. Molecular Ecology, 2018. 27(19): p. 3799–3810.

55. Lyon, R.L., A useful secondary sex character in Dendroctonus bark beetles. The Canadian Entomologist, 1958. 90(10): p. 582–584.

56. Sim, S.B., et al., HiFiAdapterFilt, a memory efficient read processing pipeline, prevents occurrence of adapter sequence in PacBio HiFi reads and their negative impacts on genome assembly. BMC Genomics, 2022. 23(1): p. 157.

57. Cheng, H., et al., Haplotype-resolved de novo assembly using phased assembly graphs with hifiasm. Nature Methods, 2021. 18(2): p. 170–175.

58. Li, H. and R. Durbin, Fast and accurate short read alignment with Burrows-Wheeler transform. Bioinformatics, 2009. 25(14): p. 1754–60.

59. Li, H., et al., The Sequence Alignment/Map format and SAMtools. Bioinformatics, 2009. 25(16): p. 2078–2079.

60. Zhou, C., S.A. McCarthy, and R. Durbin, YaHS: yet another Hi-C scaffolding tool. Bioinformatics, 2022. 39(1).

61. Durand, N.C., et al., Juicer provides a one-click system for analyzing loop-resolution Hi-C experiments. Cell Syst, 2016. 3(1): p. 95–8.

62. Durand, N.C., et al., Juicebox provides a visualization system for Hi-C contact maps with unlimited zoom. Cell Syst, 2016. 3(1): p. 99–101.

63. Simão, F.A., et al., BUSCO: assessing genome assembly and annotation completeness with single-copy orthologs. Bioinformatics, 2015. 31(19): p. 3210–3212.

64. Flynn, J.M., et al., RepeatModeler2 for automated genomic discovery of transposable element families. Proceedings of the National Academy of Sciences, 2020. 117(17): p. 9451–9457.

65. Gel, B. and E. Serra, karyoploteR: an R/Bioconductor package to plot customizable genomes displaying arbitrary data. Bioinformatics, 2017. 33(19): p. 3088–3090.

66. Zhbannikov, I.Y., et al., SeqyClean: A pipeline for high-throughput sequence data preprocessing, in Proceedings of the 8th ACM International Conference on Bioinformatics, Computational Biology,and Health Informatics. 2017, Association for Computing Machinery: Boston, Massachusetts, USA. p. 407–416.

67. Kim, D., et al., Graph-based genome alignment and genotyping with HISAT2 and HISAT-genotype. Nature Biotechnology, 2019. 37(8): p. 907–915.

68. Stanke, M., et al., Gene prediction in eukaryotes with a generalized hidden Markov model that uses hints from external sources. BMC Bioinformatics, 2006. 7(1): p. 62.

69. Stanke, M. and S. Waack, Gene prediction with a hidden Markov model and a new intron submodel. Bioinformatics, 2003. 19 **Suppl 2**: p. ii215–25.

70. Porubsky, D., et al., SVbyEye: a visual tool to characterize structural variation among whole-genome assemblies. Bioinformatics, 2025. 41(6).

71. Godefroid, M., et al., Restriction-site associated DNA markers provide new insights into the evolutionary history of the bark beetle genus Dendroctonus. Molecular Phylogenetics and Evolution, 2019. 139: p. 106528.

72. Loytynoja, A., Phylogeny-aware alignment with PRANK. Methods Mol Biol, 2014. 1079: p. 155–70.

73. Zhang, Z., KaKs_Calculator 3.0: Calculating selective pressure on coding and non-coding sequences. Genomics, Proteomics & Bioinformatics, 2022. 20(3): p. 536–540.

74. Schrider, D.R., et al., Rates and genomic consequences of spontaneous mutational events in Drosophila melanogaster. Genetics, 2013. 194(4): p. 937–954.

75. Keightley, P.D., et al., Estimation of the spontaneous mutation rate in Heliconius melpomene. Mol Biol Evol, 2015. 32(1): p. 239–43.

76. Yang, S., et al., Parent–progeny sequencing indicates higher mutation rates in heterozygotes. Nature, 2015. 523(7561): p. 463–467.

77. Six, D.L. and R. Bracewell, Chapter 8 - Dendroctonus, in Bark Beetles, F.E. Vega and R.W. Hofstetter, Editors. 2015, Academic Press: San Diego. p. 305–350.

78. Yan, L., ggvenn: draw Venn diagram by ‘ggplot2’. 2021, R package.

79. Quinlan, A.R., BEDTools: The Swiss-Army Tool for Genome Feature Analysis. Curr Protoc Bioinformatics, 2014. **47**: p. 11 12 1–34.

80. Ranwez, V., et al., MACSE v2: toolkit for the alignment of coding sequences accounting for frameshifts and stop codons. Mol Biol Evol, 2018. 35(10): p. 2582–2584.

81. Dainat, J., et al., NBISweden/AGAT: AGAT-v1.4.1. 2024, Zenodo.

82. Team, R.C., R: A Language and Environment for Statistical Computing. 2025.

83. Buchfink, B., K. Reuter, and H.G. Drost, Sensitive protein alignments at tree-of-life scale using DIAMOND. Nat Methods, 2021. 18(4): p. 366–368.

84. Cantalapiedra, C.P., et al., eggNOG-mapper v2: functional annotation, orthology assignments, and domain prediction at the metagenomic scale. Mol Biol Evol, 2021. 38(12): p. 5825–5829.

85. Pertea, M., et al., StringTie enables improved reconstruction of a transcriptome from RNA-seq reads. Nat Biotechnol, 2015. 33(3): p. 290–5.

