## Supplemental Figs for "The tortured past of young polymorphic sex chromosomes revealed through multiple *de novo* genome assemblies of the mountain pine beetle"

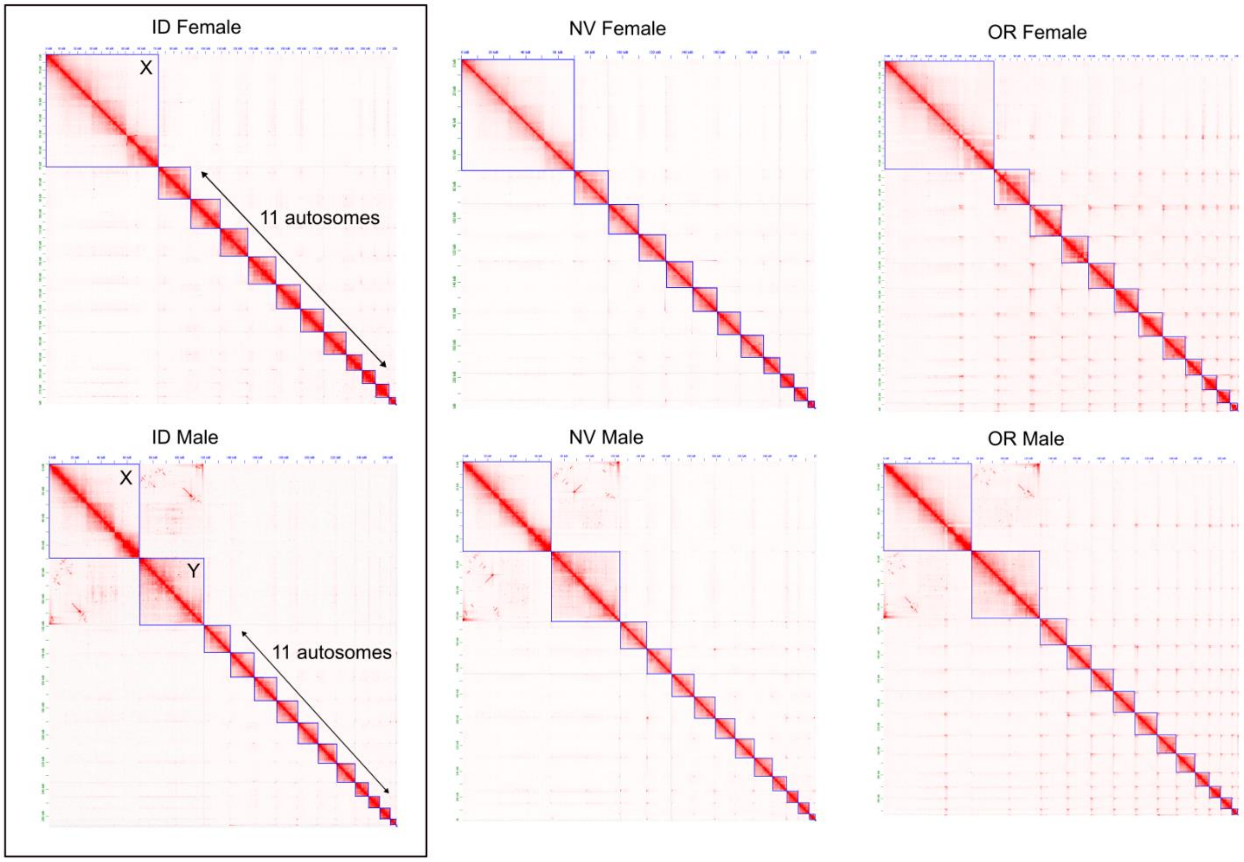


**Supplemental Figure 1. Hi-C heatmaps of all six MPB genome assemblies.** Blue boxes denote chromosomes. Note that all chromosomes are oriented in the same direction in all heatmaps with putative centromere shown to the left for acrocentric chromosomes.

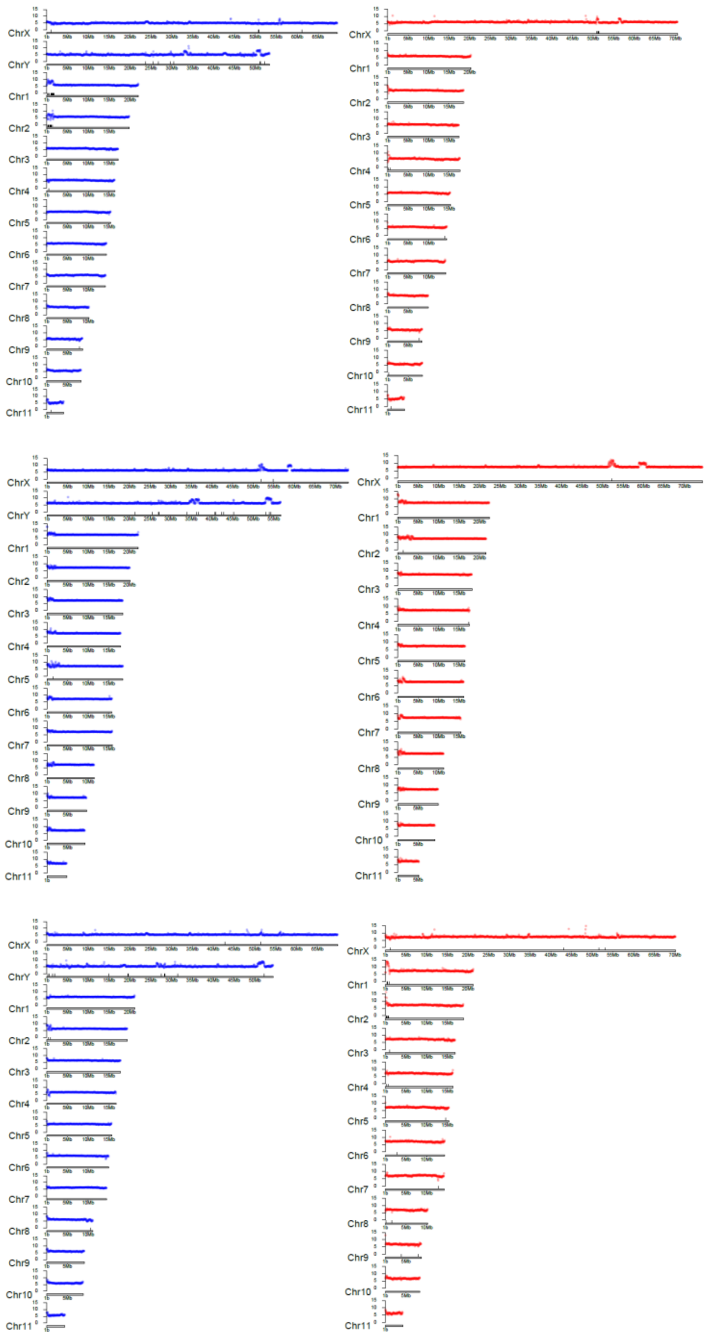

**Supplemental Figure 2. PacBio HiFi sequencing coverage.** ID male (top left) and female (top right), OR male (middle left) and female (middle right), and NV male (bottom left) and female (bottom right) assemblies. Log2 coverage in 50 kb windows (blue = male assemblies, red = female assemblies) and scaffolding stitch points between contigs (tick marks) are shown.

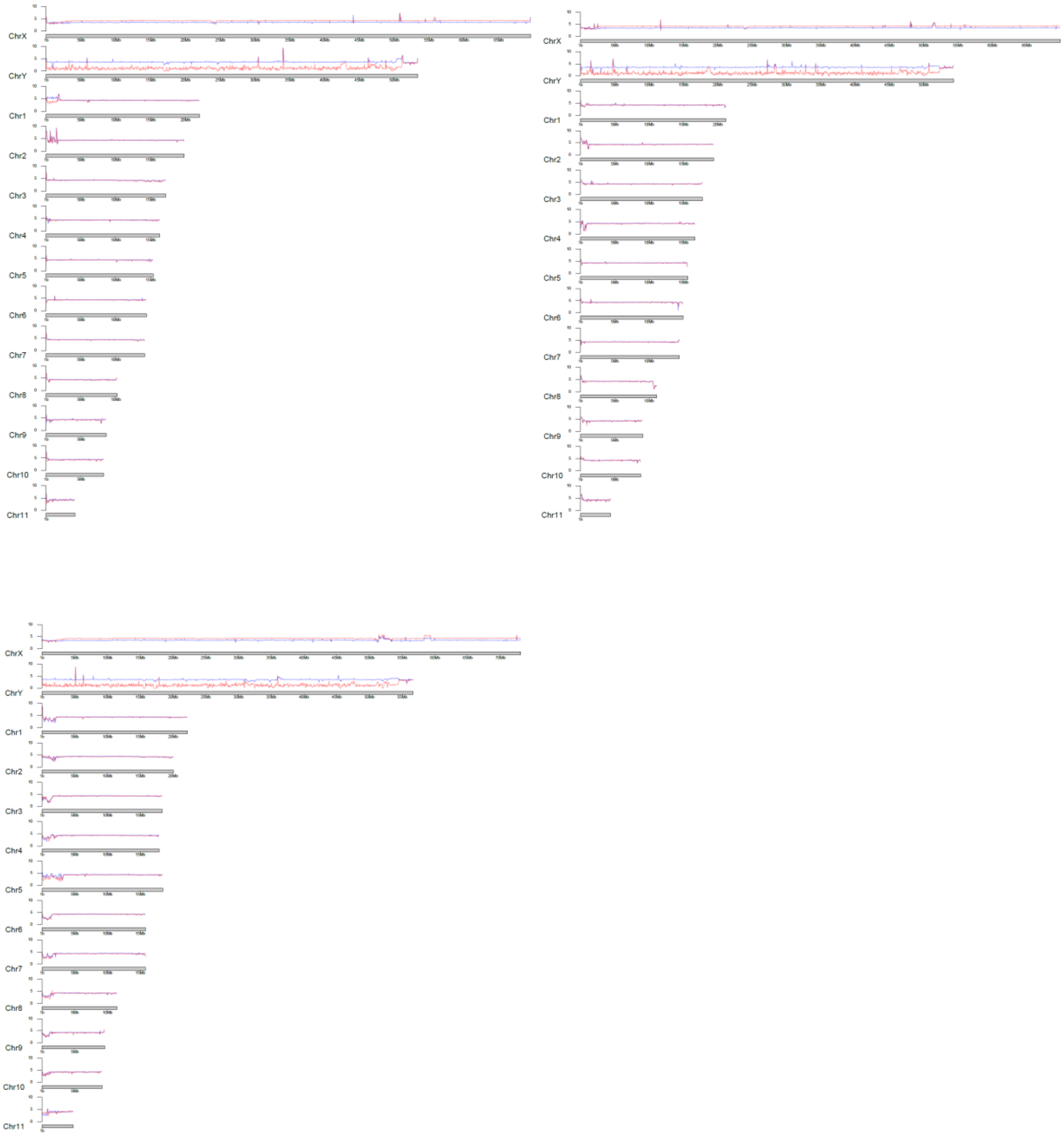


**Supplemental Figure 3. Illumina sequencing coverage of genome assemblies.** Log2 male (blue) and female (red) Illumina sequencing coverage of the ID male (top left), NV male (top right), and OR male (bottom left) reference genome assemblies.

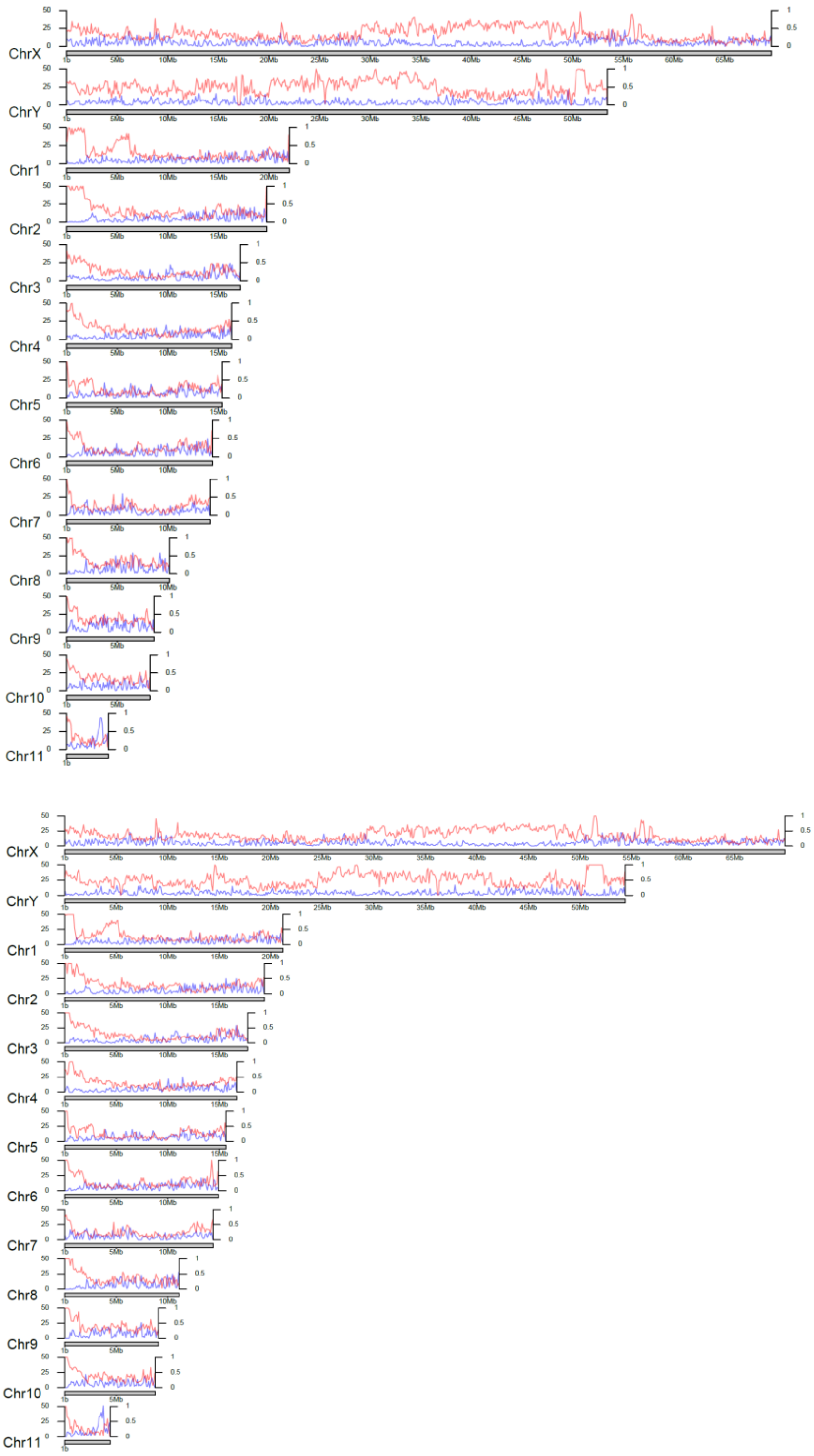


**Supplemental Figure 4. Repeat and gene density.** Number of protein coding genes (blue, left axis) and proportion of bases repeatmasked (red, right axis) estimated in 50 kb windows in ID (top), and NV (bottom) male genome assemblies.

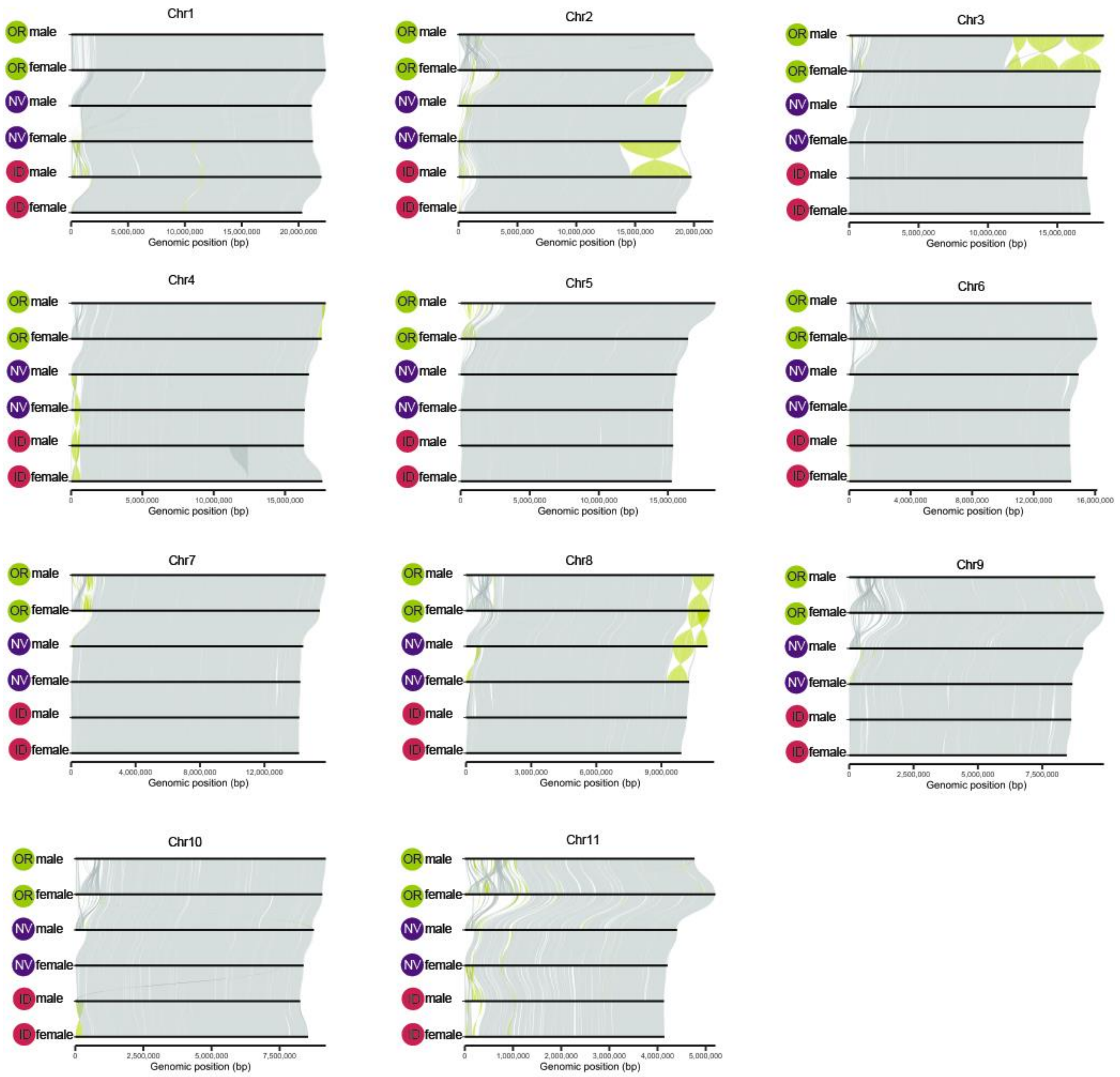


**Supplemental Figure 5**. **Whole-genome alignments of autosomes for all six MPB genome assemblies.** Inferred acrocentric centromeric regions are shown to the left for all chromosomes. Inverted regions are highlighted in yellow. Note we oriented all autosomes to have the putative pericentromeric region at the start.

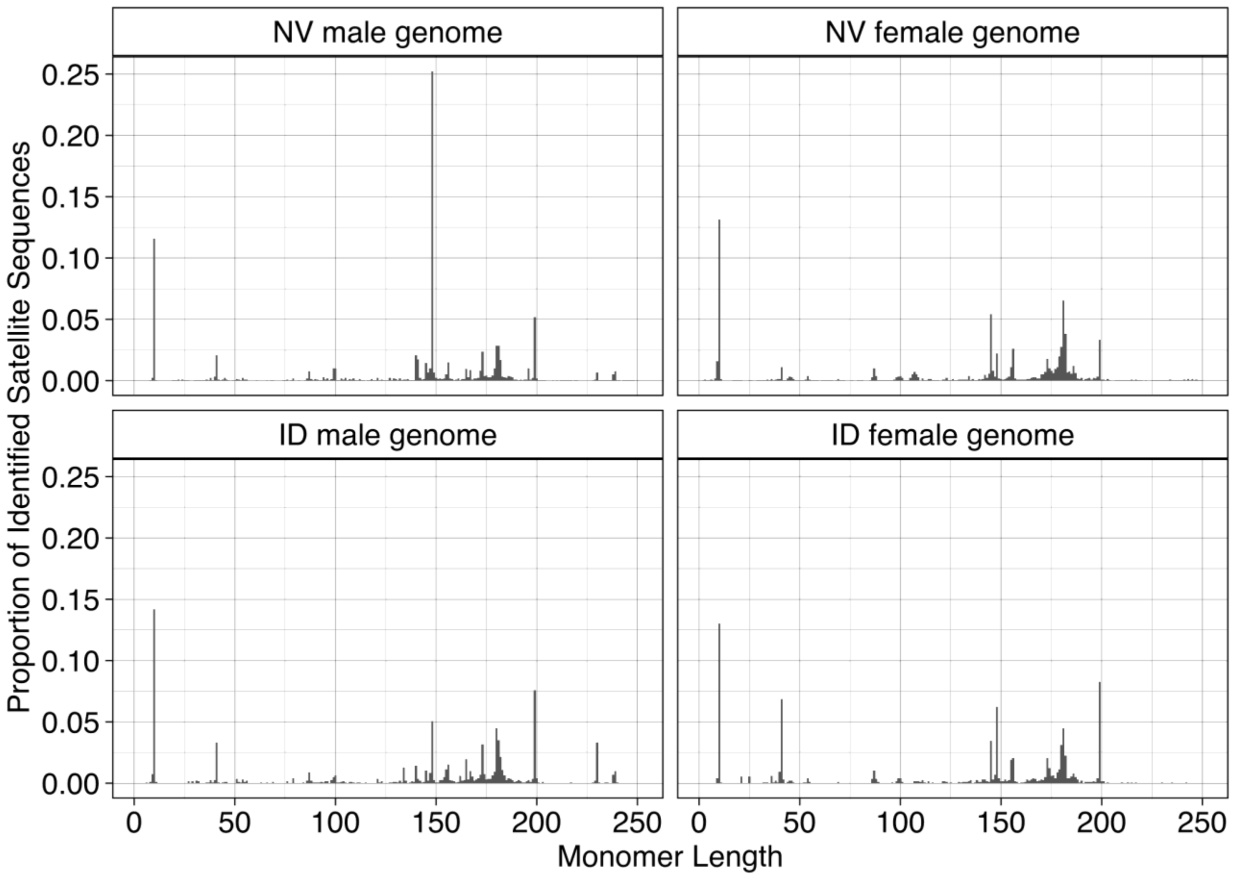


**Supplemental Figure 6. The distribution of identified satellite sequences identified by TRASH arranged by monomer length in each genome.** The monomers at 10 (putative telomere) and 148 and 199 (putative centromere-associated sequence) consistently stand out as monomer peaks in all assemblies. We are only visualizing monomers from 0-250.

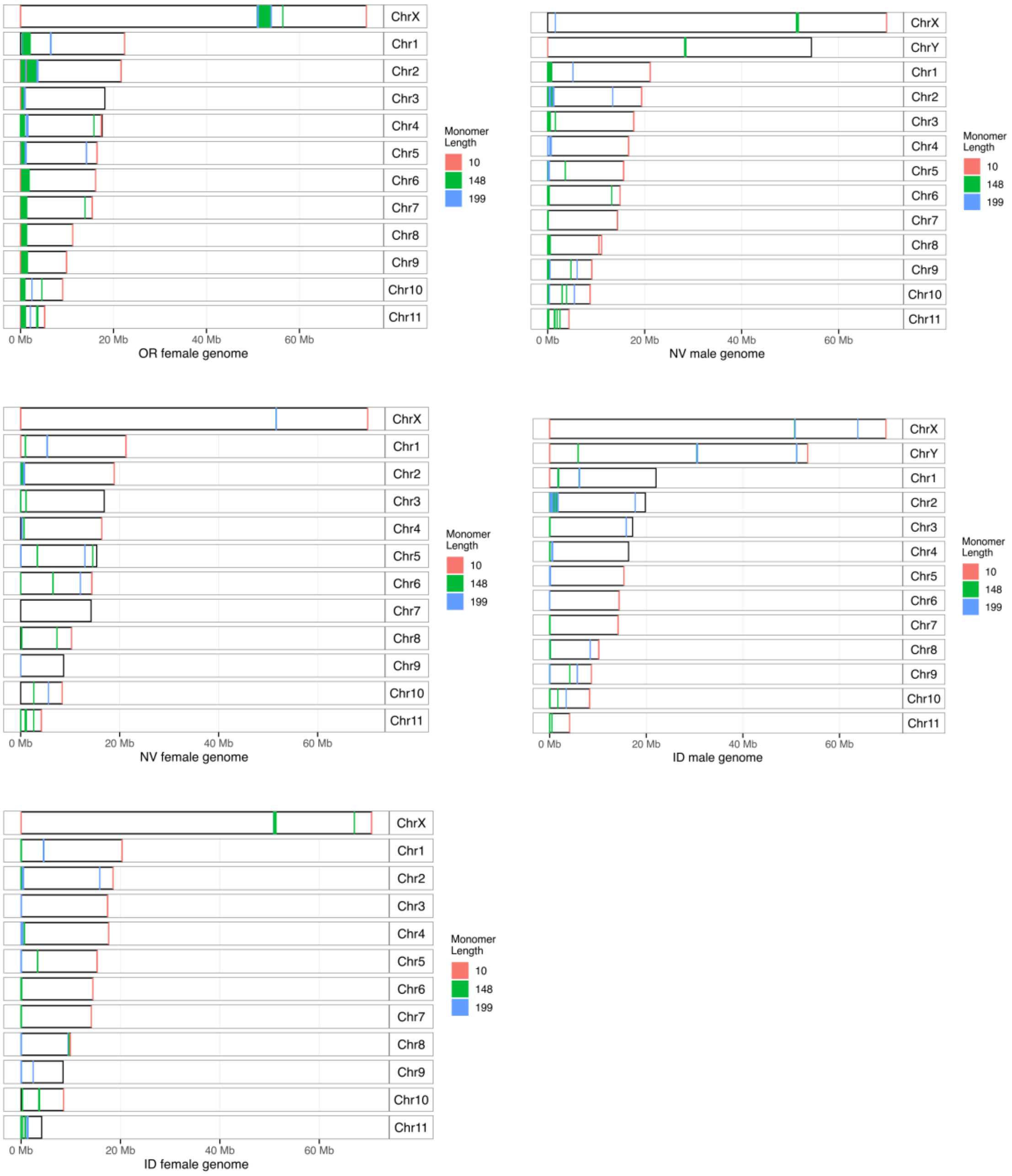


**Supplemental Figure 7. Genomic distribution of centromere and telomere-associated satellite sequence across chromosomes**. The 10-mer is putatively associated with telomere and the 148- and 199-mers are putatively associated with the centromere. Autosomes are inferred acrocentric while the X and Y are both metacentric.

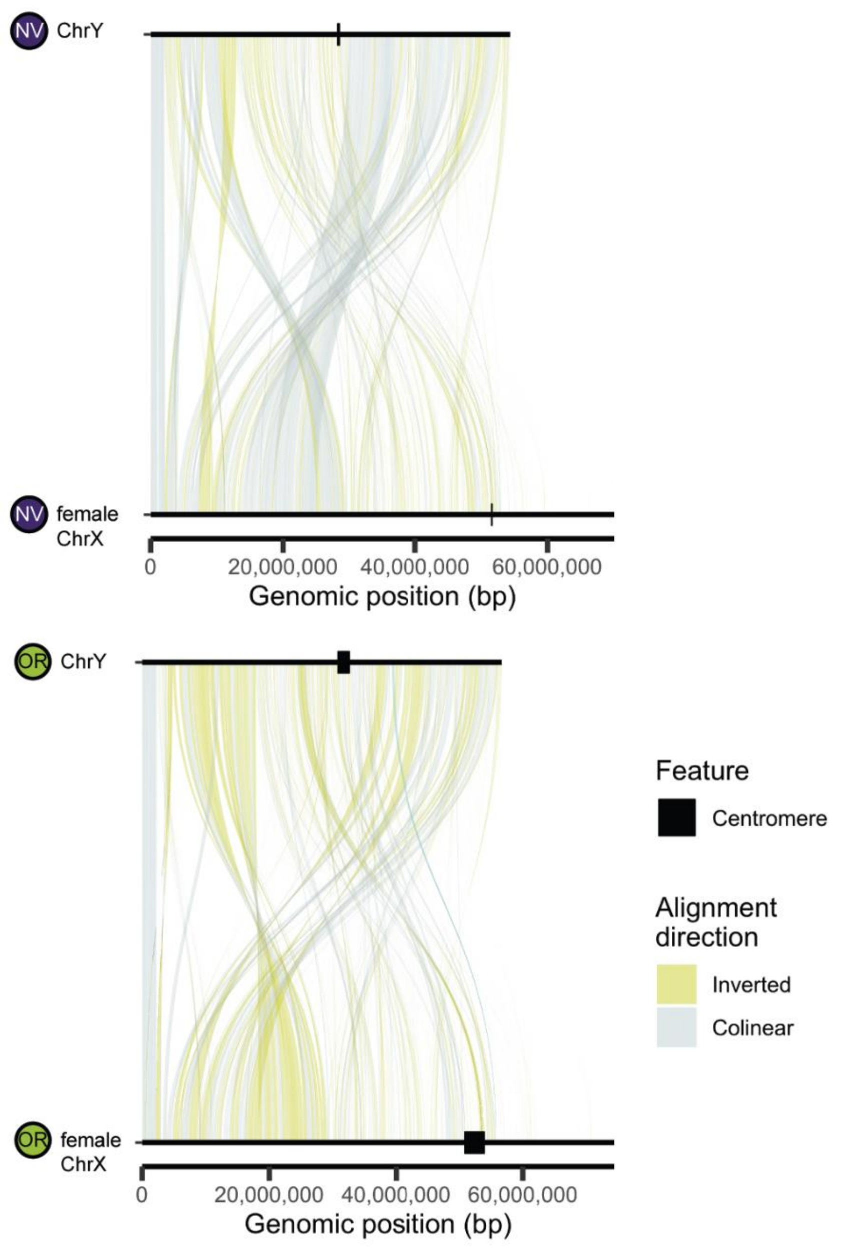


**Supplemental Figure 8. Chromosome alignments between neo-X and neo-Y reveal numerous rearrangements.** Colinear regions shown in gray and inverted regions are highlighted in yellow. The centromeres are denoted in black. The neo-Ys are reversed in orientation to align the PARs at 0. We identified 958 and 1240 inverted regions in NV and OR, respectively.

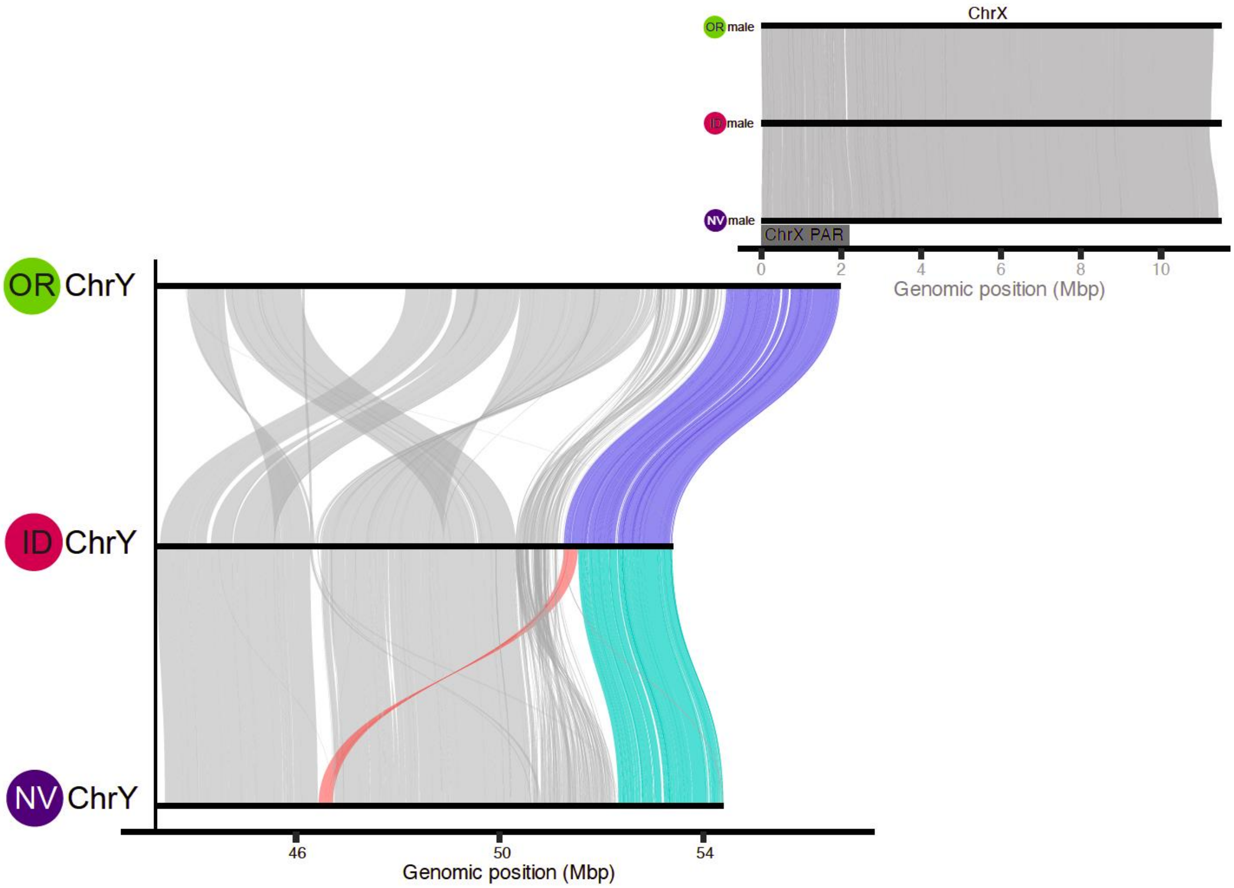


**Supplemental Figure 9**. **NV ChrY PAR is reduced in length (bp) compared to OR and ID ChrY PAR**. Chromosome alignments between OR, ID, and NV PAR and adjacent non-PAR Y. Inferred OR/ID (blue) and NV (cyan) PAR are highlighted. A portion of the OR/ID PAR has changed physical position in NV (red), likely as the result of complex inversions which have occurred repeatedly across the Y chromosome. Note the X PAR for all males is colinear (top right).

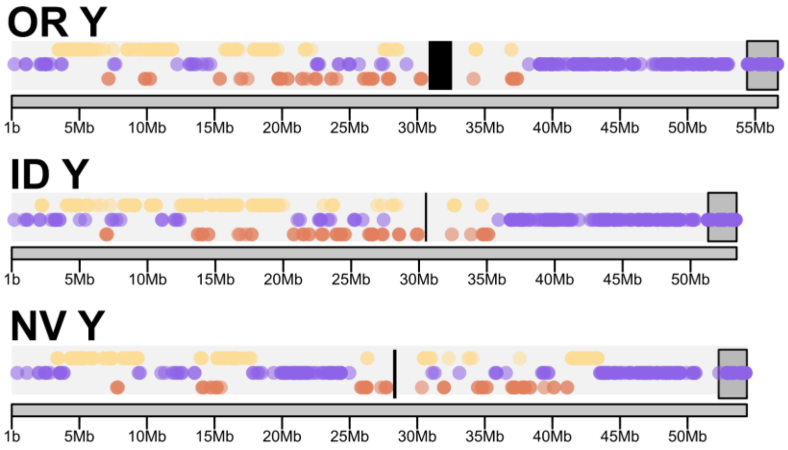


**Supplemental Figure 10. The neo-Y genomic coordinates of gametologs with a *D. valens* ortholog.** Brown points (bottom row) are orthologs from the oldest fusion (valens-Chr1) to anc-X. Yellow (valens-Chr4, top row) and purple (valens-Chr11, middle row) points are orthologs from the subsequent fusions. Grey boxes and black boxes show PARs and putative centromeres, respectively. Note the neo-X location of the same gametologs is shown in Figure 3B.

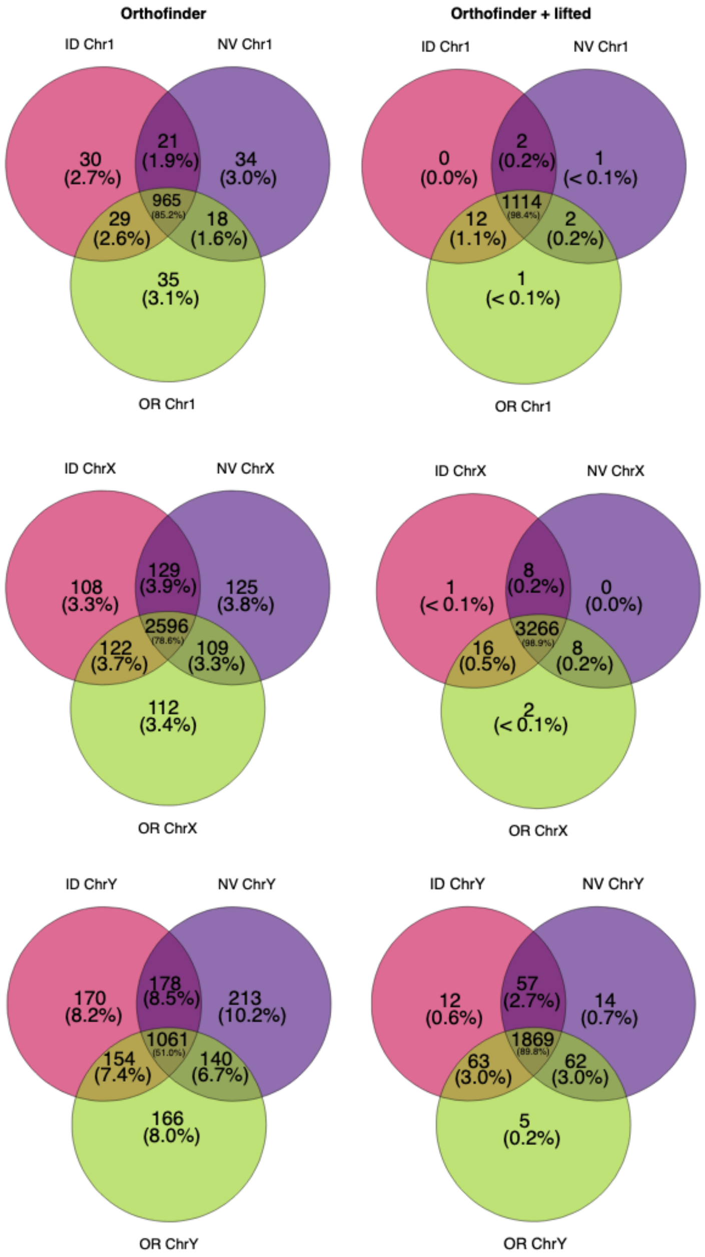


**Supplemental Figure 11. Gene counts in males for Chr1, ChrX, and ChrY with Orthofinder compared to Orthofinder with lifting genes that were initially identified as 1:1:0 or 1:0:0.**

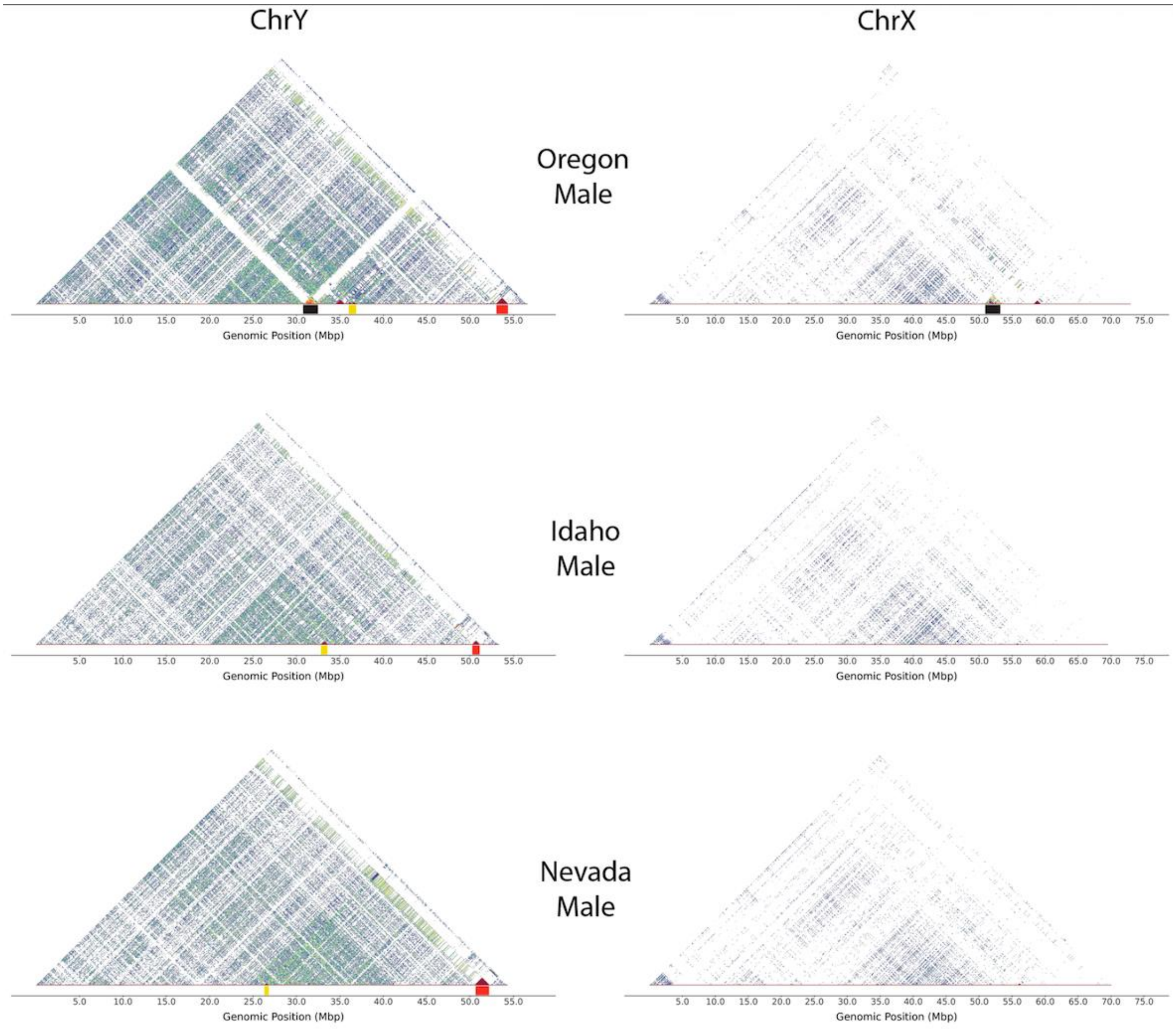


**Supplemental Figure 12. Self-identity heat maps for ChrY and ChrX in OR, ID, and NV males.** The heatmap is based on sequence identity and the range is designated as cold (blue; 86% percent identity) to hot (maroon; 100% percent identity). Centromere is notable on OR ChrX (~52 Mb, black bar) and ChrY (~31 Mb, black bar), but not ID and NV. On ChrY, red patches include a *FOXF1* gene expansion (yellow bar) and a highly repetitive region near the PAR on ChrY (~50-54Mb, red bar).

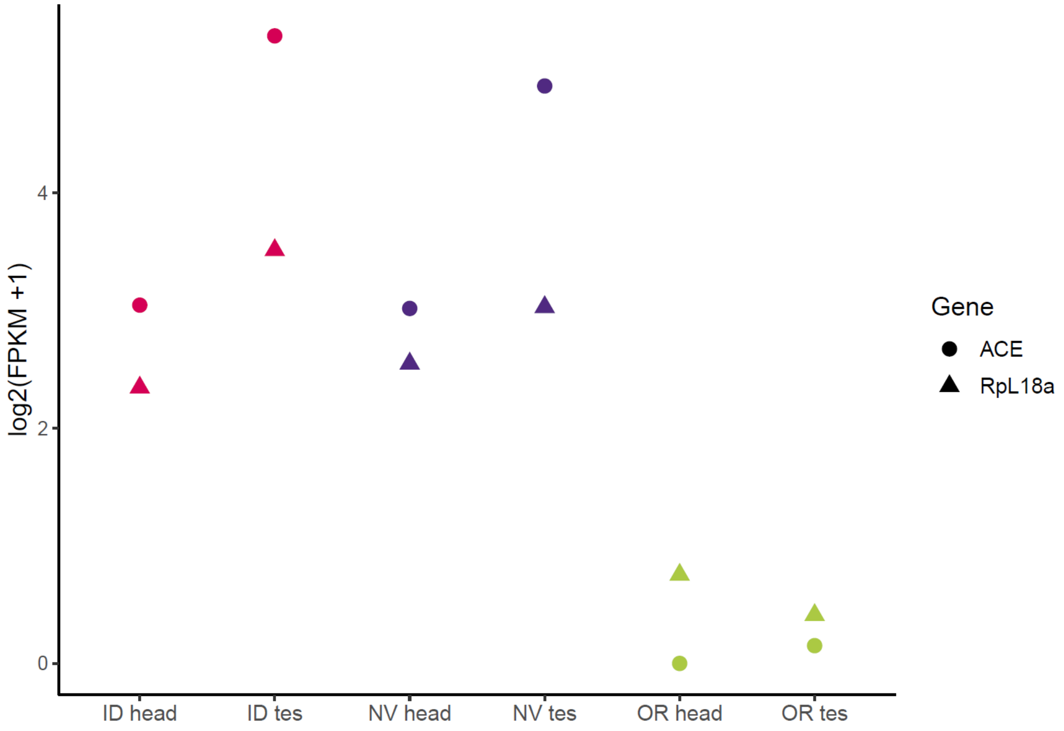


**Supplemental Figure 13. Median head and testes gene expression for *ACE* and *RpL18a*.**
